# Kinetic networks identify Twist2 as a key regulatory node in adipogenesis

**DOI:** 10.1101/2021.11.17.469040

**Authors:** Arun B. Dutta, Daniel S. Lank, Róża K. Przanowska, Piotr Przanowski, Lixin Wang, Bao Nguyen, Ninad M. Walavalkar, Fabiana M. Duarte, Michael J. Guertin

## Abstract

Adipocytes contribute to metabolic disorders such as obesity, diabetes, and atherosclerosis. Prior characterizations of the transcriptional network driving adipogenesis overlook transiently acting transcription factors (TFs), genes, and regulatory elements that are essential for proper differentiation. Moreover, traditional gene regulatory networks provide neither mechanistic details about individual RE-gene relationships nor temporal information needed to define a regulatory hierarchy that prioritizes key regulatory factors. To address these shortcomings, we integrate kinetic chromatin accessibility (ATAC-seq) and nascent transcription (PRO-seq) data to generate temporally resolved networks that describe TF binding events and resultant effects on target gene expression. Our data indicate which TF families cooperate with and antagonize each other to regulate adipogenesis. Compartment modeling of RNA polymerase density quantifies how individual TFs mechanistically contribute to distinct steps in transcription. Glucocorticoid receptor activates transcription by inducing RNA polymerase pause release while SP and AP1 factors affect RNA polymerase initiation. We identify *Twist2* as a previously unappreciated effector of adipocyte differentiation. We find that TWIST2 acts as a negative regulator of 3T3-L1 and primary preadipocyte differentiation. We confirm that *Twist2* knockout mice have compromised lipid storage within subcutaneous and brown adipose tissue. Previous phenotyping of *Twist2* knockout mice and Setleis syndrome (*Twist2*^*-/-*^) patients noted deficiencies in subcutaneous adipose tissue. This network inference framework is a powerful and general approach for interpreting complex biological phenomena and can be applied to a wide range of cellular processes.

## Introduction

Mature adipocytes contribute to a multitude of metabolic processes by regulating energy balance, producing hormones, and providing structural and mechanical support (Rosen and Spiegelman 2006). Adipocyte hyperplasia downstream of increased adipogenesis is associated with pathogenesis of obesity, type 2 diabetes, and cardiovascular disease (Unamuno et al. 2018; Van Kruijsdijk et al. 2009). Adipogenic factors represent opportunities for intervention and possible mitigation of obesity-related sequelae (Ahmad et al. 2020; Ghaben and Scherer 2019). Adipocyte maturation is a tightly regulated process involving many chromatin and transcriptional changes downstream of TF binding (Madsen et al. 2020; Rauch et al. 2019; Siersbæk et al. 2011; Thompson et al. 2016; Tsankov et al. 2015). While prior studies have extensively characterized the TFs and gene expression changes required for adipogenesis (Lefterova and Lazar 2009; Rosen and MacDougald 2006; Siersbæk et al. 2012), this work relied on measurements taken hours or days apart on cells undergoing adipogenesis. Molecular events, such as TF binding, chromatin remodeling, and redistribution of RNA polymerase, occur on a time scale of seconds to minutes (Chen et al. 2014; Duarte et al. 2016; McNally et al. 2000). Therefore, previous examinations of adipogenic signaling likely omitted multiple waves of signaling and potential regulatory factors that may be critical to the process.

Molecular genomics assays can query transcriptional events with extremely high temporal resolution. While each assay delivers a tremendous amount of information, each is limited in the biology that it measures. ChIP-seq directly quantifies chromatin occupancy of proteins, but the assay is dependent upon availability of antibodies and limited to a single factor at a time. ATAC-seq and DNase-seq assays quantify chromatin accessibility, which is an indirect measure of regulatory element activity (Boyle et al. 2008; Buenrostro et al. 2015). Combining accessibility data with TF motif analyses can accurately infer TF binding without the need for factor and species-specific antibodies (Vierstra et al. 2020; Wu et al. 1979). Kinetic experiments can further increase the sensitivity of inferring dynamic TF binding, since changes in TF binding modulate local chromatin structure and accessibility (Guertin and Lis 2010, 2013; Siersbæk et al. 2011, 2014). However, these assays do not directly inform on changes in transcription and RNA polymerase dynamics. While RNA-seq is a popular approach for measuring transcription, the assay relies on accumulation of mature RNA species over hours, making it inappropriate for rapid measurements. In addition, it is difficult to deconvolve mechanistic insights from RNA-seq data, which measures secondary and compensatory transcription as well as long-lived RNA species predating initial measurements. Alternatively, nascent transcription profiling with PRO-seq captures RNA polymerase density genome-wide at high spatial and temporal resolution (Kwak et al. 2013). PRO-seq, like RNA-seq, is limited in its ability to identify potential upstream regulatory elements (REs) and regulatory transcription factors. Only by combining multiple approaches can one fully capture the signaling dynamics driving transcription regulatory cascades.

Differential TF activity defines cell identity and drives cellular responses to environmental stimuli by enforcing gene regulatory programs (Takahashi and Yamanaka 2006). Sequence-specific TFs bind to conserved motifs (Ptashne 1967) in REs within promoters and enhancers to regulate different mechanistic steps in transcription (Fuda et al. 2009). TFs recruit cofactors such as chromatin remodelers, acetyltransferases, methyltransferases, and general transcription machinery to REs. TFs are generally characterized as activators or repressors based upon their interaction partners, and recent studies more specifically describe TFs based upon their molecular function and which mechanistic steps they regulate (Danko et al. 2013; Duarte et al. 2016; Hah et al. 2011; Neumayr et al. 2022; Sathyan et al. 2019; Scholes et al. 2017). For example, pioneer transcription factors specialize in chromatin opening (Zaret and Carroll 2011). In addition to chromatin opening and RNA polymerase recruitment, many transcription steps are precisely regulated, such as RNA polymerase pausing, elongation, and termination. RNA Polymerase II (PolII) pauses *∼* 30-50 base pairs downstream of the transcription start site (TSS) (Rasmussen and Lis 1993; Rougvie and Lis 1988) and the vast majority of genes exhibit promoter-proximal PolII pausing (Core et al. 2008; Muse et al. 2007; Zeitlinger et al. 2007). Further modifications to the PolII complex triggers pause release and productive elongation (Marshall and Price 1995). Defining the steps regulated by TFs is necessary to understand how TFs coordinate with one another productively or antagonistically to regulate complex gene expression programs.

Transcriptional networks consist of multiple rapid waves of signaling through time with potential regulatory feedback and signal propagation through activation and repression of regulatory factors. These complex regulatory cascades are not captured in traditional gene regulatory networks. Differentiating one wave from the next requires observations at multiple, closely spaced time points. In this study, we perform ATAC-seq and PRO-seq on 3T3-L1 cells at seven time points within the first four hours of adipogenesis. We incorporate accessibility and transcription changes into a multi-wave signaling network and identify TF families driving the regulatory cascade. We identified Twist2 as a highly inter-connected node within the adipogenesis network. Perturbation of TWIST2 in *in vitro, ex vivo*, and *in vivo* validated the regulatory role of TWIST2 in adipogenesis. This novel network framework also infers individual regulatory relationships among TFs, REs, and target genes. The network infers TF binding events and potential mechanistic interactions between specific REs and genes. The network is designed so that one can easily identify key regulatory TF or RE hubs, determine a set of target genes for specific TFs, assess TF cooperativity, and develop testable hypotheses. We find that different transcription factor families regulate distinct mechanistic steps in the transcription cycle and coordinate with one another to orchestrate the adipogenic transcriptional cascade.

## Results

### TFs from at least 14 families are associated with dynamic chromatin accessibility in 3T3-L1 differentiation

TFs bind promoters and enhancers to modify chromatin structure and influence transcription of nearby genes. To identify dynamic REs and potential TFs that regulate adipogenic differentiation, we induced adipogenesis in 3T3-L1 mouse preadipocytes (see Methods), harvested samples at 8 time points, and performed genome-wide chromatin accessibility assays (ATAC-seq) (Figure 1A). Chromatin accessibility is a molecular measurement used to infer TF binding and RE activity. We identified over 230,000 accessibility peaks and differentiation time is the major driver of variation among the samples (Figure S1A). Approximately 13% of all peaks change significantly over the time course (Figure S1B). We clustered dynamic peaks based on kinetic profiles (Figure S1C), which resulted in five general response classes (Figure 1B). To identify candidate sequence-specific TFs that drive RE dynamics, we performed *de novo* motif analysis on dynamic peaks (Bailey et al. 2015). This approach yielded 14 potential TF family motifs including CEBP, TWIST, SP, KLF, AP1, and the steroid hormone receptor motif (Figure 1C & S1D). TF families comprise multiple proteins containing paralogous DNA binding domains that recognize very similar sequence motifs (Figure 1C). For example, multiple factors including androgen receptor, mineralocorticoid receptor, progesterone receptor, and glucocorticoid receptor (GR) bind to the steroid hormone receptor motif. However, GR is the only factor gene that is expressed in 3T3-L1 cells (Figure S1E). Therefore, we refer to the steroid hormone receptor binding consensus sequence as the GR motif. We identified AP1, CEBP, and GR, which are known positive effectors of adipogenesis (Distel et al. 1987; Flodby et al. 1996; Freytag et al. 1994; Moitra et al. 1998; Ramji and Foka 2002; Rubin et al. 1978; Siersbæk et al. 2011; Steger et al. 2010; Tanaka et al. 1997; Wang et al. 1995; Yeh et al. 1995). Members of the KLF and SP families are known to be associated with both pro-adipogenic (Birsoy et al. 2008; Inuzuka et al. 1999; Li et al. 2005; Mori et al. 2005; Oishi et al. 2005; Pei et al. 2011) and anti-adipogenic functions (Banerjee et al. 2003; Kawamura et al. 2006; Sue et al. 2008; Tang et al. 1999). The TWIST family of TFs have previously unappreciated roles in adipogenesis, but have been shown to be important for differentiation of other mesenchymal cell types, such as osteoblasts (Bialek et al. 2004; Yousfi et al. 2001). Members of all these factor families are expressed in 3T3-L1 cells (Figure S1E).

**Fig. 1.**
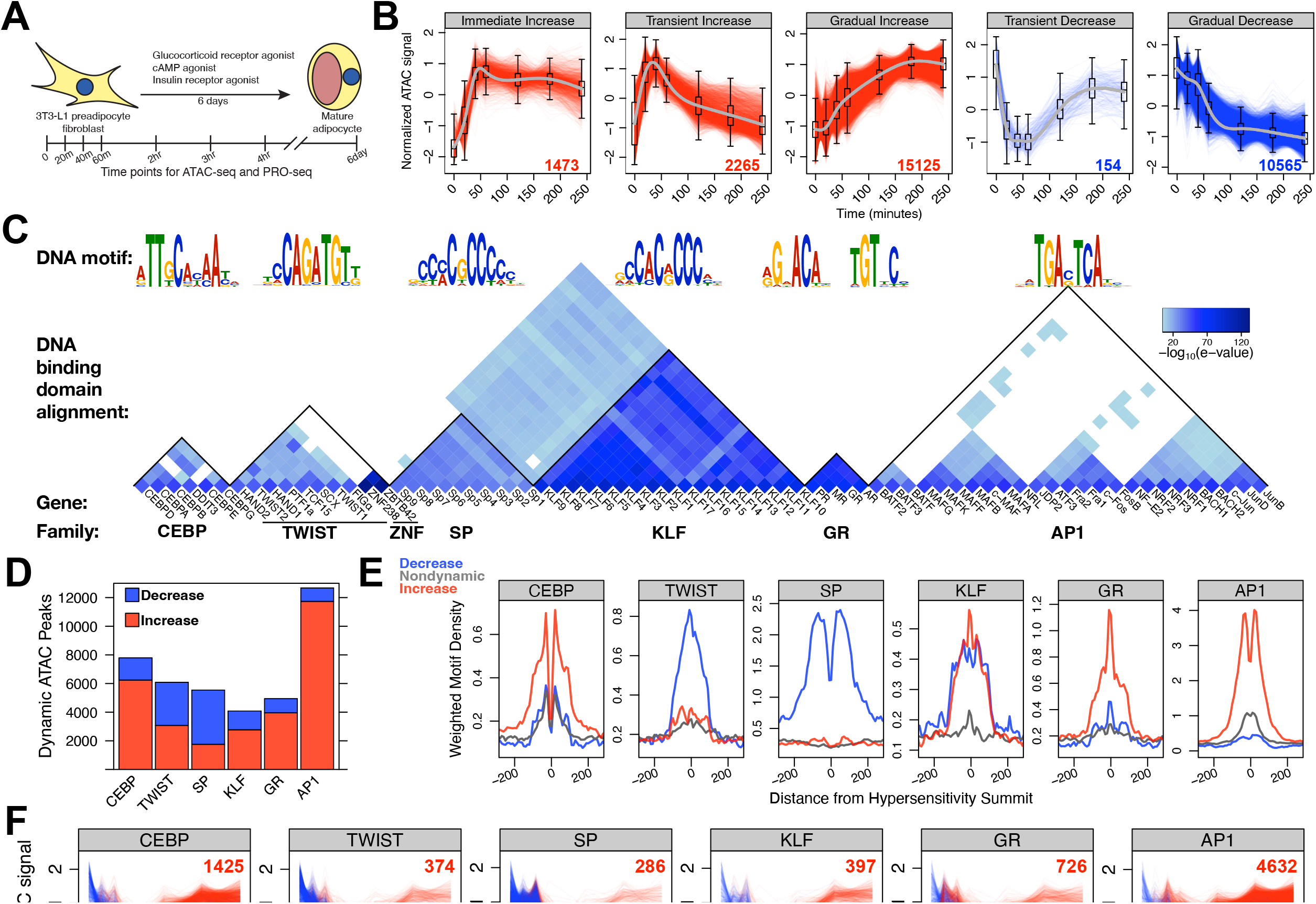
CEBP, TWIST, SP, KLF, GR, and AP1 TF families drive either increased or decreased chromatin accessibility in adipogenesis. A) Preadipocyte fibroblast 3T3-L1 cells were treated with an adipogenesis cocktail and harvested at the indicated time points for ATAC-seq and PRO-seq experiments. B) Temporal classification of ATAC-seq peaks revealed five major dynamic classes. Each dynamic ATAC peak is a red or blue trace with the number of peaks in the class indicated in the lower right; the x-axis represents time and the y-axis indicates normalized accessibility. C) *De novo* motif analysis identified the top six DNA motifs enriched within dynamic peaks. The individual TFs listed in the wedge below the seqLogo recognize the respective DNA motifs. The heatmap quantifies the local protein sequence alignment of the DNA binding domains for the genes, as determined by the Smith-Waterman algorithm (Farrar 2007). D) Dynamic ATAC-seq peaks are classified by the presence of each DNA motif. The red bars represent the number of dynamic ATAC-seq peaks within the Immediate Increase, Transient Increase, and Gradual Increase categories; the blue bars correspond to the Transient Decrease and Gradual Decrease classes. E) Red, blue, and grey traces are composite motif densities relative to ATAC peak summits for the increased, decreased, and nondynamic peak classes. The y-axis quantifies the density of the indicated position-specific weight matrix and each motif instance is weighted by its conformity to a composite motif. F) Dynamic traces of peaks that exclusively contain the specified motif indicate that CEBP, GR, and AP1 associate with increasing accessibility; SP and TWIST associate with decreasing accessibility. Peak traces are colored as in panel (B). These conclusions are consistent with the reciprocal analysis from panel (E).

TF binding or dissociation from DNA leads to enrichment of cognate motifs in dynamic peaks. The biological functions of the TFs determine whether binding or dissociation results in increased or decreased accessibility. Binding of TFs that recruit activating cofactors, such as histone acetyltransferses or remodeling enzymes that eject nucleosomes, can increase accessibility; dissociation of these factors decreases chromatin accessibility. Likewise, binding and dissociation of factors that recruit deacetylases, repressive methyltransferases, or DNA methyltransferases can affect accessibility. We found that the majority of peaks containing CEBP, KLF, GR, or AP1 motifs increase accessibility, while peaks containing TWIST or SP motifs decrease accessibility (Figure 1D). We performed the reciprocal analysis and plotted the density of motif instances relative to the summits of increased, decreased, and nondynamic peak classes to confirm the classification (Figure 1E & S1F). AP1, GR, and CEBP motifs are strongly enriched around summits of increased peaks, while TWIST and SP motifs are enriched around summits of decreased peaks. SP and KLF families have paralogous DNA binding domains and recognize similar motif sequences; however, we confidently associate chromatin decondensation to KLF factors and chromatin condensation to SP factors (Figure S1G). The SP family is composed of canonical activators (McKnight and Kingsbury 1982), therefore SP TFs are likely dissociating from the chromatin to reduce accessibility. Although we ascribe opening and closing functions to the KLF and SP families, it is impossible to determine the relative contribution of KLF and SP factors at any individual motif. We believe that the dual enrichment of KLF motifs at both increased and decreased peak summits is due to erroneous classification of SP-bound REs as KLF-bound REs. This complication is not limited to closely related motifs, as many dynamic peaks contain multiple factor binding motifs, making it difficult to isolate the contribution of individual factors. To address this complication, we plotted the changes in accessibility at dynamic peaks that contain only a single motif (Figure 1F). This confirmed that the majority of isolated AP1, GR, CEBP, and KLF motif-containing peaks increase in accessibility, while TWIST and SP motif-containing peaks decrease. The biological interpretation of these results is that the adipogenic cocktail activates members of the AP1, GR,

CEBP, and KLF TF families both directly and through transcriptional activation of family member genes, leading to RE binding and chromatin decondensation. SP and TWIST motifs are associated with decreased accessibility. TFs can act as repressors by binding to chromatin and recruiting chromatin modifiers such as deacetylases. Alternatively, dissociation of an activating TF can lead to gene repression. These results confirm the importance of several TF families and suggest that previously unappreciated TF families, such as TWIST, contribute to adipogenesis.

### SP, NRF, E2F6, KLF, and AP1 factor motifs are associated with bidirectional transcription at regulatory elements

Coordinate TF binding ultimately results in the recruitment of RNA polymerases and initiation of transcription. In mammals, core promoters and enhancers often lack sequence information that consistently orient initiating RNA polymerases (Core et al. 2014). Therefore we sought to identify bidirectional transcription signatures as a complement to chromatin accessibility assays to identify REs (Core et al. 2008; Danko et al. 2013; Seila et al. 2008). We captured the short lived divergent transcripts found at active REs with PRO-seq in parallel with the ATAC-seq adipogenesis time points (Figure 1A). We used discriminative regulatory-element detection (dREG) to identify peaks of bidirectional transcription from our PRO-seq data (Wang et al. 2019). We identified over 180,000 dREG peaks (Figure 2A & B) and 18% change significantly over the time course (Figure S2A). ATAC-seq and PRO-seq measure distinct but related biological phenomenon, therefore they identify different but overlapping sets of REs. Approximately 22% of dynamic dREG peaks overlap with dynamic ATAC-seq peaks, compared to 20% of dynamic ATAC-seq peaks in the inverse comparison. To further analyze the two classes of REs, we separated the dynamic dREG and ATACseq peaks into intragenic, intergenic, and promoter regions (Figure 2C). Both methods effectively identify REs within promoters (Figure S2B). We find PRO-seq more sensitively detects intragenic REs relative to the other categories, while ATAC-seq efficiently detects intergenic REs. We closely evaluated the overlap between ATAC and dREG peaks by plotting PRO-seq signal at ATAC peaks and vice versa (Figure S2C). We observe the distinctive bidirectional transcription signature at ATAC peaks irrespective of whether or not the ATAC peaks intersect dREG peaks. The signature is less intense at ATAC-seq peaks that do not overlap dREG peaks. Likewise, ATAC-seq signal is enriched at dREG-peaks that do not overlap ATAC-seq peaks (Figure S2D). Moreover, dREG peaks within intergenic, intragenic, or promoter regions that do not overlap with ATAC-seq peaks have less bidirectional transcription (Figure S2E). Although we find that bidirectional transcription and accessibility do not perfectly correlate, we are likely underestimating the extent of accessibility and bidirectional transcription overlap.

**Fig. 2.**
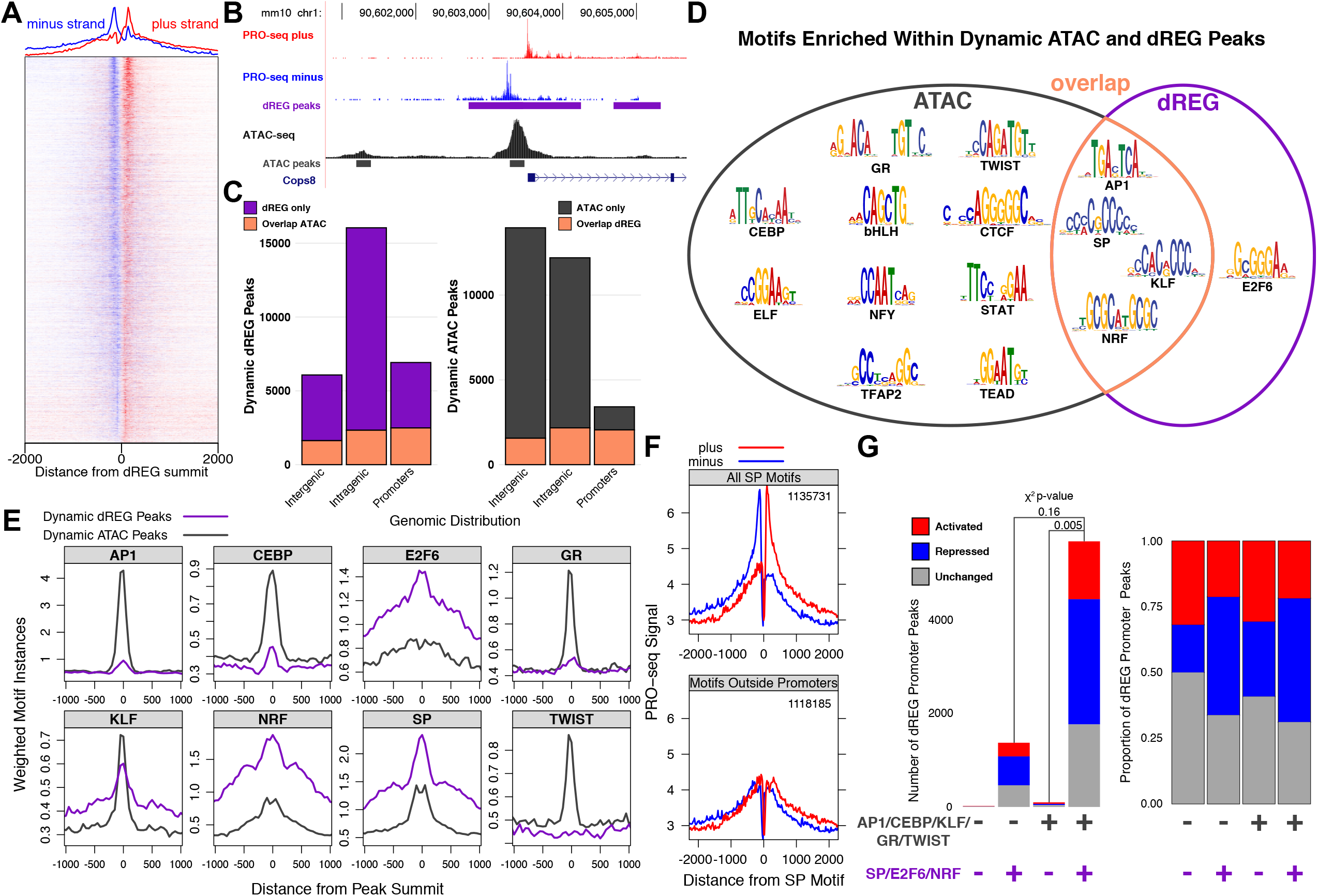
SP, NRF, and E2F6 TF families drive bidirectional transcription dynamics at regulatory regions within gene bodies and promoters. A) The heatmap illustrates over 200,000 putative REs with a bidirectional transcription signature. B) Both dREG and ATAC-seq identify a RE within the promoter of *Cops8*. The intragenic RE is only identified by its bidirectional PRO-seq signature while the upstream intergenic RE is only identified by ATAC-seq. C) Dynamic ATAC-seq and dREG-defined REs largely overlap in promoter regions. Intragenic regions are defined based on primary transcript annotation of PRO-seq data, promoters are between 150 bp upstream and 50 bp downstrean of TSSs, and intergenic regions are the remainder of the genome. D) Dynamic ATAC-seq peaks are enriched for a more diverse set of TF motifs than dynamic dREG peaks. E) Motif density distinguishes TFs associated with dynamic bidirectional transcription from those associated with dynamic accessibility. For example, TWIST and GR motifs are enriched within dynamic ATAC-seq peaks but are rarely found within dynamic dREG peaks. F) SP is only associated with bidirectional transcription at promoters and not distal REs. The top plot shows the average normalized PRO-seq signal for plus and minus strands around all 1,135,731 SP motif instances while the bottom plot displays all SP motifs excluding those in promoters (1,118,185). The distinct dual peak profile of bidirectional transcription collapses when only considering SP motifs outside promoters. G) Dynamic bidirectional transcription peaks found in promoters are stratified by the presence or absence of TF motifs. The left plot quantifies the total number of peaks and the right plot scales to the proportion of peaks in each category. The x-axis factor motif categories are defined by the presence or absence of ATAC-associated factors (AP1, CEBP, GR, KLF, and TWIST) and dREG-associated factors (SP, E2F6, and NRF). dREG-associated factor motifs are enriched in peaks that decrease bidirectional transcription, suggesting a link between SP, NRF, and E2F6 factors and an early and pervasive decrease in promoter initiation at their target genes.

We sought to identify TFs that drive bidirectional transcription by further characterizing PRO-seq-defined REs. We hypothesized that different sets of TF motifs are enriched within REs defined by ATAC-seq and PRO-seq. For instance, the cognate motifs of TFs that recruit initiation machinery may be preferentially enriched at dREG-defined REs. We performed *de novo* motif analysis on dynamic dREG peaks and found enrichment of AP1, SP, KLF, NRF, and E2F6 motifs (Figure 2D). Of these, only the E2F6 motif was not also enriched in ATAC-seq peaks. We plotted motif density around the summits of either dynamic ATAC or dREG peaks to further differentiate ATAC and dREG-defined REs (Figure 2E & Figure S2F). Of the motifs found *de novo* in dREG peaks, only E2F6, NRF, and SP were more enriched in dynamic dREG versus dynamic ATAC-seq peaks. We hypothesize that these three factor families regulate bidirectional transcription in adipogenesis. The SP motif is found in over 25% of human and mouse promoters, making the SP motif the most enriched cis-regulatory element (RE) within promoters (Benner et al. 2013). To determine whether divergent transcription signatures found at SP motifs are dominated by SP factors within promoters, we plotted plus and minus strand nascent transcription at all SP motif instances (Figure 2F top). Indeed, when SP motifs within promoters are removed from the composite input, divergent transcription peaks collapse (Figure 2F bottom). We also observe this phenomenon with E2F6 and NRF motifs (Figure S2G), implying that these factors and SP preferentially regulate divergent transcription at promoters. Next, we wanted to determine whether SP, NRF, and E2F6 motifs within promoters associate with increasing or decreasing divergent transcription. We find that bidirectional transcription tends to decrease in REs with dREG-enriched motifs as opposed to those without dREG-enriched motifs (Figure 2G). This further supports the previous conclusions that SP and NRF motifs associate with decreases in RE activity (Figure 1E & Figure S1G). Although ATAC-seq and PRO-seq are both orthogonal methods to identify putative REs, we find that ATAC-seq is more sensitive at identifying distal regulatory TFs and PRO-seq primarily captures functional TFs within promoters.

### Defining predicted TF binding events as *trans*-edges in the network

We determined candidate functional TFs within the set of REs by searching for over-represented sequence motifs and determining the expression levels of TF family members. However, inferring TF binding from accessibility, motif, and expression data at any individual site remains a challenge (Guertin et al. 2012; Li et al. 2019). In addition to chromatin accessibility, expression of the TF, and presence of the TF’s cognate motif, we leverage the change in accessibility over the time course to infer TF binding and dissociation events in adipogenesis. We term these predicted changes in TF occupancy, which are directed linkages from TFs to REs, as *trans*-edges in our networks. For simplicity, we refer to *trans*-edges as factor binding or dissociation events.

We define the following rules for *trans*-edge inference: 1) The RE must first be defined as an ATAC-seq peak at any time point. 2) The binding motif of the upstream TF must be present in RE. 3) Chromatin accessibility must change significantly between two time points to infer binding or dissociation. 4) The direction of accessibility changes must match with the molecular function of the TF as defined in Figure 1. 5) Members of the TF family must be expressed at the appropriate time point; for example, the TF must be expressed at the later time point for binding and the earlier time point for dissociation. 6) GR, AP1, and CEBP are directly activated by the adipogenic cocktail, so we infer edges from expressed family members to REs from 0-20 minutes. We necessitate that the nascent RNA expression of the other TFs changes significantly to infer *trans* edges from their genic node to an RE node. Mechanistically, TFs have short residency times on DNA and they are continually binding and dissociating from their sites *in vivo* (Chen et al. 2014; McNally et al. 2000). When we refer to inferred binding and dissociation within the network, we are strictly referring to overall changes in occupancy at a genomic site within the population of cells.

The following examples highlight implementations of these rules. The *Nr3c1* gene, which encodes GR, decreases expression immediately upon treatment (Figure S3A). We suggest that the rapid transcriptional repression of *Nr3c1* is the reason GR-associated increases in accessibility are transient. Therefore, we restrict binding edges attributed to GR to the first 40 minutes of the time course. We attribute any significant decreases in accessibility at inferred GR binding REs observed at later time points to dissociation of GR. We label these edges with a *dissociation* attribute. In the case of SP, we find that *Sp1, Sp3*, and *Sp4* are all repressed early in the time course (Figure S3B). We hypothesize that the delayed accessibility decrease associated with SP motifs is due to transcriptional repression and natural turnover of the SP pool, which results in overall dissociation of SP on chromatin (Figure 1F). We restrict *trans*-edges for SP to the later part of the time course. Conversely, we observe *Twist2* gene activation early in the time course (Figure S3C). Therefore, we predict that TWIST-associated repression is a result of increased TWIST binding and recruitment of negative co-factors. *Twist2* expression levels have returned to baseline in mature adipocytes (Figure S3D), suggesting that TWIST’s effects are transient and can only be captured with an early, high-resolution time course. By focusing only on the REs that change accessibility and integrating with transcription data, we infer TF binding and dissociation events that drive adipogenesis.

We integrated publicly available TF ChIP-seq datasets to assess the performance of *trans*-edge inference. Specifically, we incorporated ChIP-seq profiling of AP1 (cJun and JunB), KLF (KLF4 and KLF5), CEBP*—*, and GR (Siersbæk et al. 2011, 2014). All these experiments were performed in 3T3-L1 preadipocytes at 4 hours of differentiation. We found that 60-70% of the inferred binding events for these factors overlap called ChIP-seq peaks (Figure S3E). The one exception was for GR, which exhibited a much lower degree of overlap (35%). This is consistent with our network, which suggests that GR binds and dissociates rapidly from the chromatin at many sites. ChIP-seq experiments performed at 4 hours may be too late to capture these transient binding events. Furthermore, CEBP, GR, and KLF regulatory elements that exhibit sustained high accessibility (i.e. nonattenuated) displayed a higher degree of overlap with ChIP-seq peaks than those that did not, suggesting that our ATAC-seq dynamics captured fluctuations in factor binding. In addition, we plotted composite ChIP signal at our predicted binding sites and found a strong enrichment in signal at REs that overlap with ChIP-seq peaks (Figure S3F). We also observed a weaker enrichment of ChIP signal around inferred binding events that do not overlap with ChIP-seq peaks, suggesting weaker binding events at these locations was overlooked in ChIP-seq peak calling. Direct measurement of factor binding by ChIP-seq validates the predictive power of using dynamic ATAC signal and the presence of sequence motifs to infer factor binding.

### Proximal changes in accessibility are tightly linked to transcription

Chromatin accessibility positively correlates with local gene transcription. We confirmed this assertion by quantifying transcription of genes within 10 kilobases (kb) of dynamic ATAC-seq peak sets that exclusively increase or decrease accessibility (Figure 3A). The majority of genes (63%) with one proximal increasing ATAC-seq peak are activated, likewise 68% of genes proximal to a single decreasing ATAC-seq peak are repressed. Genes near two or more increased accessibility peaks are much more likely to be associated with transcription activation, and vice versa (Figure 3A). To further validate this association and explore the relationship between RE and target gene distance, we focused on all genes near one dynamic peak and stratified gene/peak pairs based on distance between the TSS and peak summit (Figure S3G-I). The closer the peak and the gene, the more likely gene transcription and peak accessibility correlate in the same direction. This result indicates that proximal REs have a greater impact on gene expression than distal elements. Moreover, we plotted change in gene transcription against distance-scaled local accessibility changes and observed the expected positive correlation between transcription and accessibility at both early and late phases of the time course (Figure S3J & K). These findings indicate that both accessibility dynamics and distance are important factors when considering the relationship between REs and genes.

**Fig. 3.**
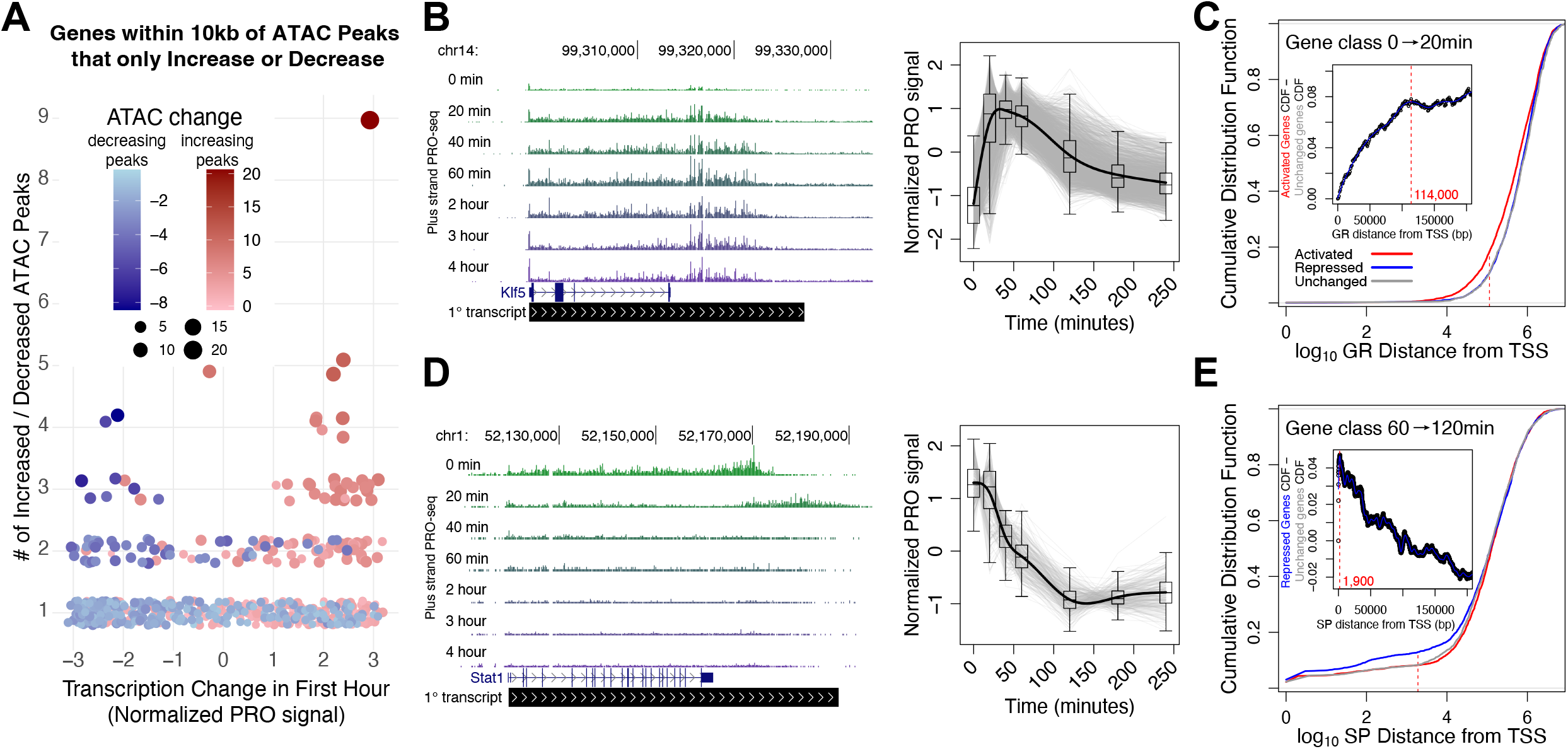
Chromatin accessibility, transcription dynamics, and proximity guide inference of *cis*-edges between REs and target genes. A) Change in gene expression correlates with local accessibility change over the first hour. Each data point represents a gene within 10 kb of either only increased (red) or decreased (blue) peaks. The y-axis indicates the number of increased or decreased accessibility peaks near the gene and the x-axis represents the normalized change in gene transcription over the first hour. B) *Klf5* (left) is part of a cluster of 1717 immediately and transiently activated genes (gray traces on the right). C) Cumulative distribution plots showing distance between GR-bound REs and genes either activated (red), repressed (blue), or unchanged (grey) over the first hour of the time course. The left-shift of the red curve suggests that ATAC-seq peaks with GR motifs are closer to the 20 vs. 0 min activated gene class. The inset plot reports the difference in cumulative distribution between the activated and unchanged gene classes as distance from the TSS increases. The leveling off of the traces at 114 kb from the TSSs suggests that GR-mediated transcription activation requires GR to bind within 100 kb of the TSS. D) *Stat1* (left) is part of a cluster of repressed genes (gray traces on the right). E) ATAC-seq peaks with SP motifs are closer to the 60 vs. 120 min repressed gene class.Traces converge 1,900 bases from the start sites, suggesting that the functional distance of SP-mediated gene repression is within 2 kb of TSSs.

We incorporated the distance between REs and genes as well as covariation in their accessibility and transcription to infer functional links, termed *cis*-edges, in our networks. We define *cis*-edges as predicted regulatory relationships between REs and genes. For example, if GR binds a regulatory element within a gene’s promoter and induces gene activation, we would draw a *cis*-edge between the RE and the gene. Within the network, we assign GR as an attribute to the edge. To confidently infer and annotate *cis*-edges, we must assess whether a class of TFs is associated with increasing or decreasing transcription. Since the distance between a RE and a gene influences the likelihood that accessibility and transcription will covary, we classified the function of a TF class within the context of adipogenesis by determining if peaks with a cognate TF motif are closer to activated or repressed gene classes. We expected that factor families associated with decreases in accessibility, like SP, would be closer to repressed genes on average. To test this hypothesis, we first categorized genes as significantly activated, repressed, or unchanged for each pairwise comparison within the time course. For example, *Klf5* (Figure 3B left panel) is one of a subset of 4225 genes immediately activated from 0 to 20 minutes (Figure 3B right panel). Plotting the cumulative distribution function (CDF) for genes against the distance between the closest peak summit and the gene TSS shows that GR peaks tend to be closer to the 4225 activated genes compared to the repressed or unchanged genes (Figure 3C). To estimate the maximum range that a factor can act, we plotted the difference between the repressed gene class CDF against the unchanged gene class CDF against distance between gene and peak (Figure 3C inset). We find that the difference in the CDFs plateaus at around 114 kb, meaning that inferred GR binding events accumulate at the same rate for activated and unchanged genes at distances greater than 114 kb. This distance constraint represents an empirical observation that suggests a maximum regulatory distance in this system. It is possible that we would detect a different constraint in another cellular context due to differences in genomic architecture and regulatory environment. Immediately repressed genes like *Stat1* are closer to SP peaks than control gene sets (Figure 3D & E). This analysis indicates that SP acts very proximal to its target genes, with an actionable range of less than 2 kb (Figure 3E inset). This finding is consistent with our previous conclusions that decreased SP peaks are primarily found in promoters (Figure 2F). Our observed maximal distances represent a hypothesized maximal actionable distance for factor activity in 3T3-L1s. However, we anticipate that optimal distance for most factors is much closer to target genes than these maxima. Therefore we apply a closer distance threshold when inferring regulatory relationships between individual REs and genes as described in the next section. The lower thresholds increase our confidence in our predicted *cis*-edges.

### Linking REs to target genes

We incorporate these biological principles into logical rules to define *cis*-edge predictions within an adipogenesis network. We develop our rules to maximize confidence in our predicted regulatory interactions. First, a gene and regulatory element must be within 10 kb to infer a *cis*-edge. Second, RE accessibility and gene transcription must covary over the same time range. These two logical rules provisionally link REs and genes, then we employ additional rules that reflect the biology of individual TFs. For instance, the 10 kb distance metric is made more strict for factors like SP, for which the functional distance constraint as determined in Figure 3E is less than 10 kb. Temporal rules also influence edge predictions. For instance, GR-bound REs are only significantly closer to genes activated in comparison to the 0 minute time point such as the 20 vs. 0 comparison (Figure 3C), meaning that genes activated later in the time course cannot be directly activated by GR binding in the network. Therefore, as with *trans*-edges, we only infer *cis*-edges between GR-bound REs and genes that change early in the time course. Incorporating these observations into our *cis*-edge rules we infer direct functional relationships between regulatory elements, bound TFs, and changes in target gene expression.

### Constrained networks identify genes regulated combinatorially or by individual TF families

Quantifying nascent transcription with PRO-seq maps the position and orientation of RNA polymerase with base-pair resolution. Nascent transcriptional profiling captures engaged RNA polymerase species throughout the genome, including intragenic features such as the proximal promoter and gene body. We can infer regulatory mechanisms of gene sets by quantifying relative changes in RNA polymerase density within the pause region and gene body. For instance, if the rate of RNA polymerase pause release increases between conditions, we expect that the signal in the pause region to decrease and the gene body signal to increase. Previous studies focus on biological systems where one TF dominates the response, such as ER, HSF, and NF-*κ*B (Danko et al. 2013; Duarte et al. 2016; Hah et al. 2011). In these systems, the composite RNA polymerase signals at activated genes highlight differences in densities between pause and gene body compartments (Danko et al. 2013; Duarte et al. 2016; Hah et al. 2011; Sathyan et al. 2019). A complication in our system is that multiple TFs cooperate to drive transcription changes, making it difficult to identify the target steps (i.e. initiation, pause release) that TFs regulate. In order to address this complication, we identified genes that are predominantly regulated by a single TF in our network.

We constructed a bipartite network inferring changes in TF binding (*trans* edges) that regulate downstream changes in transcription (*cis* edges). Genes and REs can be regulated or bound by either one or a combination of TFs. For example, we constructed a constrained network with RE and gene nodes downstream of individual TFs, including AP1. In this network, 1224 genes are solely activated by AP1 and 1847 genes activated by AP1 and at least one other factor (Figure 4A). Most REs downstream of AP1, both individually and combinatorially bound, are not linked to any downstream genes (12608 v. 4829). This network highlights a paradigm in the transcription field that a minority of TF binding events lead to changes in gene expression (Spradling et al. 1975; Westwood et al. 1991). We constructed similar networks for GR (Figure 4B), SP (Figure 4C), CEBP (Figure S4A), KLF (Figure S4B), and TWIST (Figure S4C). These networks illustrate the interconnectivity of gene regulation, while simultaneously identifying genes that are predominantly regulated by individual factors.

**Fig. 4.**
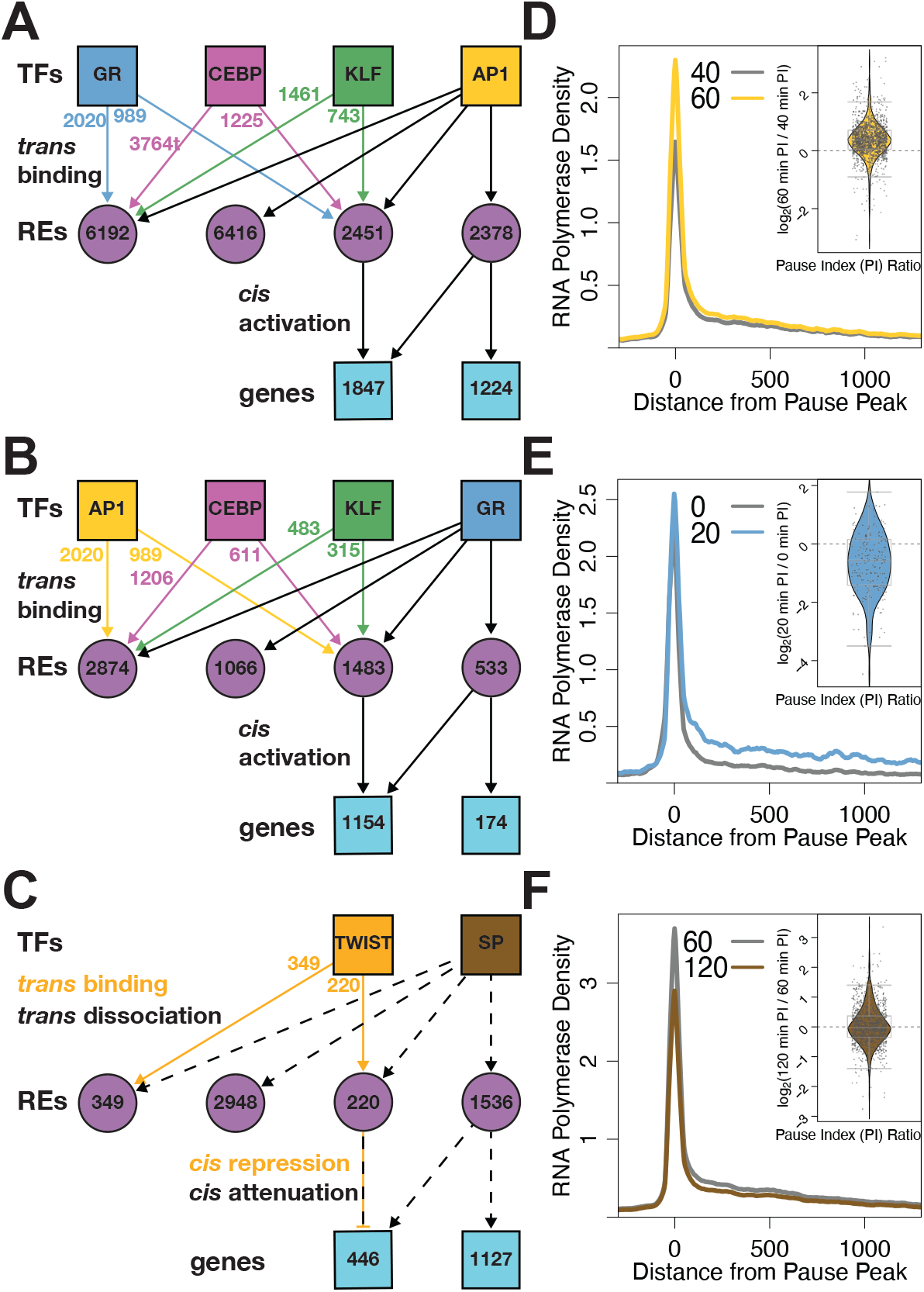
Constrained networks downstream of AP1, GR, and SP identify genes regulated by individual factors. Simplified networks highlight the number of REs and genes that are combinatorially or individually regulated by (A) AP1, (B) GR, and (C) SP. Factors bind / dissociate from REs (purple circles) and regulate genes (blue squares). Colored arrows and numbers indicate the contribution of non-lead factors to RE activity. Combinatorially regulated REs are bound by the lead TF and either one or more of the other TFs. Composite PRO-seq signal is plotted relative to the promoter-proximal pause peak of (D) 1224 genes solely regulated by AP1, (E) 174 genes regulated by GR, and (F) 1127 genes regulated by SP. Inset violin plot illustrate the change in pause index for the gene set for the indicated time points. Each data point is a gene and all genes were input from the composite.

To extract mechanistic information from genes regulated by only one TF, we plotted composite RNA polymerase density from our PRO-seq data around pause peak summits at different time points for the isolated genes (Figure 4D, E, F & Figure S4D, E, F). The resulting traces show the characteristic pause peak centered around 0 followed by release of the RNA polymerase into the gene body. Examining RNA polymerase density traces of genes only activated by AP1 at 60 and 40 minutes show an increase in density in the pause region, suggesting increased RNA polymerase recruitment to AP1-activated genes (Figure 4D). These time points were chosen because AP1 peaks are closest to genes activated between 60 and 40 minutes, suggesting that AP1 exerts the most transcriptional control during this time range. At first glance we see a similar result for GR when comparing traces from 0 and 20 minutes (Figure 4E). However the situation becomes more complex when considering the ratio of pause density to gene body density, or pause index (PI). The PI for genes regulated solely by AP1 increases on average from 40 minutes to 60 minutes (Figure 4D inset). Conversely, the PI for 71% of genes regulated solely by GR decreases on average between 0 and 20 minutes. This suggests that GR primarily activates transcription by inducing pause release (Figure 4E inset). The affected step is unique to the factor, as isolated AP1 genes don’t exhibit the decrease in pause index from 0 to 20 minutes observed with isolated GR genes (Figure S4G inset). Isolated GR genes show increases in pause index later in the time course, likely due to GR dissociation from the genome after the early phase of the time course and an associated decrease in pause release rate (Figure S4H). As for repressed genes, we find a decrease in pause peak and gene body intensity in predicted SP and TWIST target genes (Figure 4F & Figure S4F). We find that SP genes demonstrate a more symmetrical distribution of pause indices (Figure 4F inset). While it is likely that these factors affect PolII recruitment, we sought to develop a more rigorous approach to determine how changes in initiation and pause release rates can account for observed changes in PolII density.

### Modeling changes in regulatory transcription steps

We developed a mathematical model to further characterize how TFs target specific steps in the transcription cycle. Our model breaks up the gene unit into two compartments: a pause region and gene body region. PRO-seq directly measures RNA polymerase density within these regions for each gene. We define a series of differential equations to model polymerase density as measured by PRO-seq within the two compartments (Figure 5A). We establish rate constants representing different transcriptional steps, namely RNA polymerase recruitment / transcription initiation (*k*_*init*_), premature termination (*k*_*pre*_), pause release (*k*_*rel*_), and elongation (*k*_*elong*_) The values of the rate constants determine the predicted density within the two compartments. We vary the rate constants for *k*_*init*_, *k*_*pre*_, and *k*_*rel*_ over two orders of magnitude and vary *k*_*elong*_ rate from 600 to 6000 bases per minute to determine the effect on pause and gene body density and how the model compares to observed changes. We make the assumption that *k*_*elong*_ remains constant between time points. Since *k*_*pre*_ and *k*_*init*_ are opposing rates in the model, we cannot distinguish an increase in one rate from a decrease in another. To simplify the model, we keep *k*_*pre*_ constant between time points. We determine how changes in *k*_*init*_ combined with *k*_*rel*_ changes can account for the average density changes for the 174 isolated GR-regulated genes from Figure 4E. A wide range of rate parameters can describe the initial pause and gene body densities, but regardless of the initial rates, a narrow fold-change in these rates can account for the observed changes between time points (Figure 5B). We find that a ∼1.07-fold increase in recruitment / initiation and a *∼*1.50-fold increase in pause release explain the changes in compartment occupancy between 0 and 20 minutes (Figure 5B left). We calculated the absolute rate of initiation and residency time of PolII in the pause region based on the models and plotted a simulated PolII profile (Figure 5C). For this simulation, we chose the parameter set with an elongation rate closest to the established consensus rate of approximately 2500 bases per minute (Ardehali and Lis 2009; Jonkers and Lis 2015). Estimated pause residency time drops from 29 seconds to 19 seconds between 0 and 20 minutes as a result of the rate constant changes. Taking a similar approach, an *∼* 0.78-fold decrease in recruitment / initiation rate with a *∼* 0.94-fold change in pause release rate produces observed changes in PolII occupancy between 60 and 120 minutes for the 1127 isolated SP genes (Figure 5B middle). This corresponds to a initiation/ recruitment rate reduction from 15.1 to 11.9 polymerase molecules per minute (Figure 5D). If SP factors normally stimulate initiation, then mass action would explain dissociation of SP factors upon transcriptional repression of SP genes. Previous studies link SP1 to transcriptional initiation through interaction with the TFIID general TF (Gill et al. 1994). The observed changes in RNA polymerase composite profiles between 40 and 60 minutes at AP1 target genes are explained by 1.27 to 1.39-fold increases in initiation rate and 0.85 to 0.93-fold decreases in pause release rate (Figure 5B right). These relative changes in *k*_*init*_ and *k*_*rel*_ for AP1 targets do result in gene activation, but it was unexpected that the profiles are explained by a decrease in *k*_*rel*_. Since composite profiles represent the average of all included genes, it is possible that the composite represents a diverse set of genes that are regulated by different AP1 family members. We speculate that we could gain a more clear insight if we were able to deconstruct the AP1 targets and identify gene targets of specific AP1 factors. The above analyses indicate that we can deconvolve complex transcriptional networks to identify gene targets of individual TFs and determine which steps in the transcription cycle each TF preferentially regulates.

**Fig. 5.**
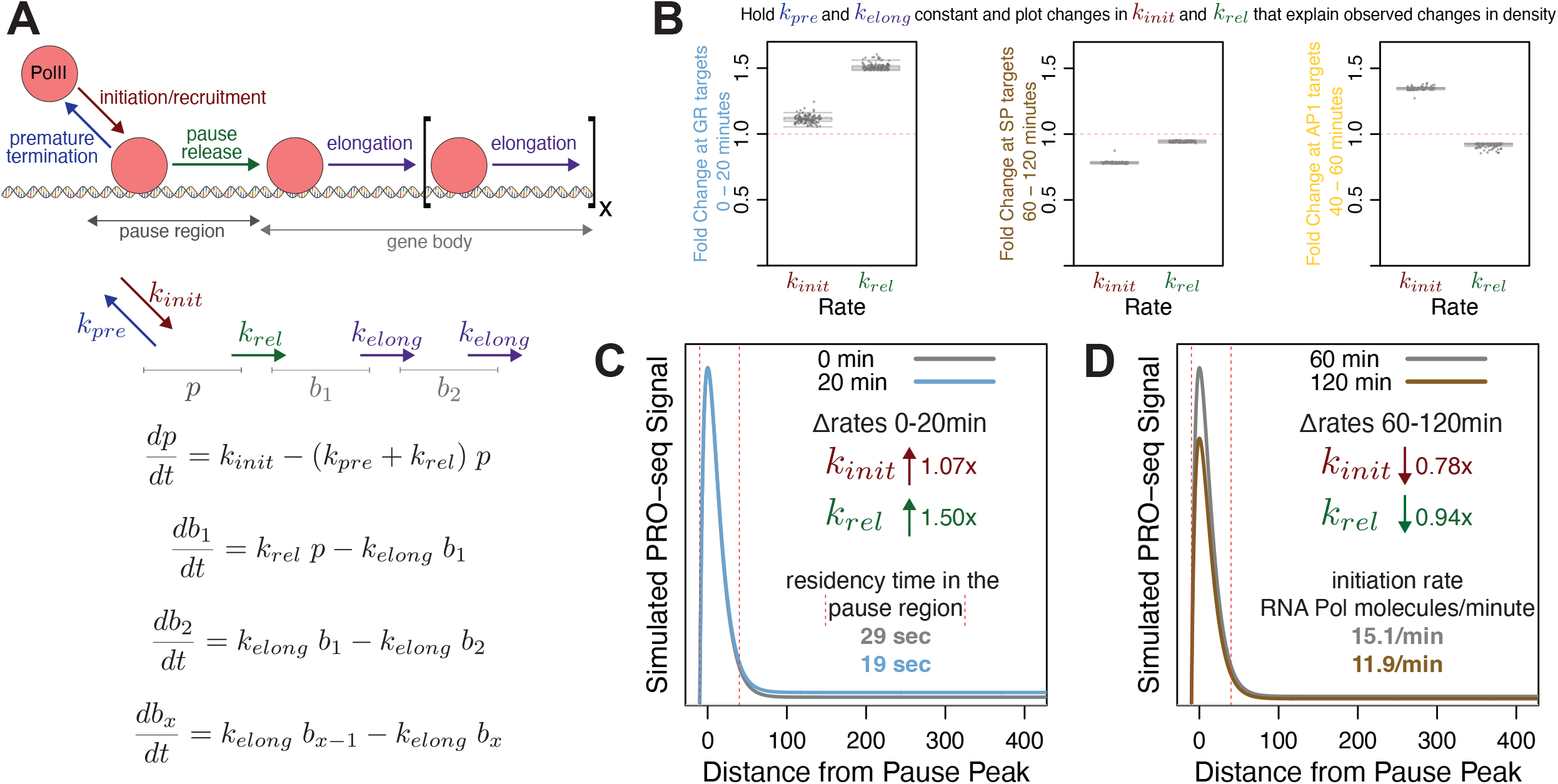
TFs induce changes to transcriptional rate constants. A) The compartment model of transcription contains rate constants for RNA Polymerase II (PolII) initiation / recruitment, premature termination, pause release, and elongation. Polymerase occupancy in the pause or gene body compartment is measured directly from PRO-seq. Differential equations relate rate constants to the rate of change of polymerase density in the compartments. B) We use the compartment model to estimate how rate constants change between 0 and 20 minutes for the 174 genes regulated solely by GR (left), 60 and 120 minutes for the 1127 genes regulated by SP (middle), and 40 and 60 minutes for the 1224 genes regulated by AP1 (right). For each estimation, we hold *k*_*pre*_ and *k*_*elong*_ constant and calculate values of *k*_*init*_ and *k*_*rel*_ that fit the observed occupancy in the pause and body regions. C-D) We simulated composite profiles for a set of parameters from (B left) and (B middle).

We applied this model and approach to a separate PRO-seq dataset of the C7 B-cell line treated with dexamethasone to determine whether GR regulates pause release within a different system in which GR is specifically activated. We identified 70 genes activated by dexamethasone treatment. The pause index of 80% of these genes decrease between 0 and 60 minutes (Figure S5A). Compartment modeling of these genes showed that a ∼1.33-fold increase in pause release rate explained the observed changes in PolII density at the activated genes (Figure S5B & C). These validation results support the role of GR regulating pause release and highlight the power of predicting the molecular function of transcription factors within complicated regulatory cascade networks generated from kinetic PRO and ATAC data.

### TFs cooperate to bind REs and activate gene expression

AP1, CEBP, GR, and KLF bind REs either individually or in combination in order to activate expression. We classify REs based on the combination of factors that bind and drive accessibility changes. Likewise, we classify genes based on which TFs are immediately upstream in the network. Genes activated by the same combination of factors can be downstream of different classes of REs. We use genes activated by all 4 of AP1, CEBP, GR, and KLF to illustrate potential regulatory scenarios. These genes may be downstream of a single RE that binds all factors (Figure 6A orange). Alternatively, the gene may be downstream of a pair of REs each binding 2 factors (Figure 6A purple), 3 and 1 (Figure 6A blue), or more complicated regulatory schemes (Figure 6A green). All 15 possible classes of REs contribute to activation of the 82 genes downstream of AP1, CEBP, GR, and KLF (Figure 6B& S6A). The largest population of RE classes is isolated AP1 peaks with 8794, while peaks bound by all activating factors is the smallest with 74. The distribution of gene classes generally mirrors the distribution of RE classes, with isolated AP1 genes being the largest class. As discussed above, all factors activate more genes in combination than in isolation. There are comparatively few combinatorially-regulated genes without AP1 contribution (1924 with AP1 v. 149 without). This finding, along with the high number of genes regulated by AP1, underscores the importance of the AP1 family in the network. While the bulk of negatively regulated genes are downstream of either SP or TWIST, approximately 20% are affected by both TWIST-mediated repression and SP-mediated attenuation (Figure S6B).

**Fig. 6.**
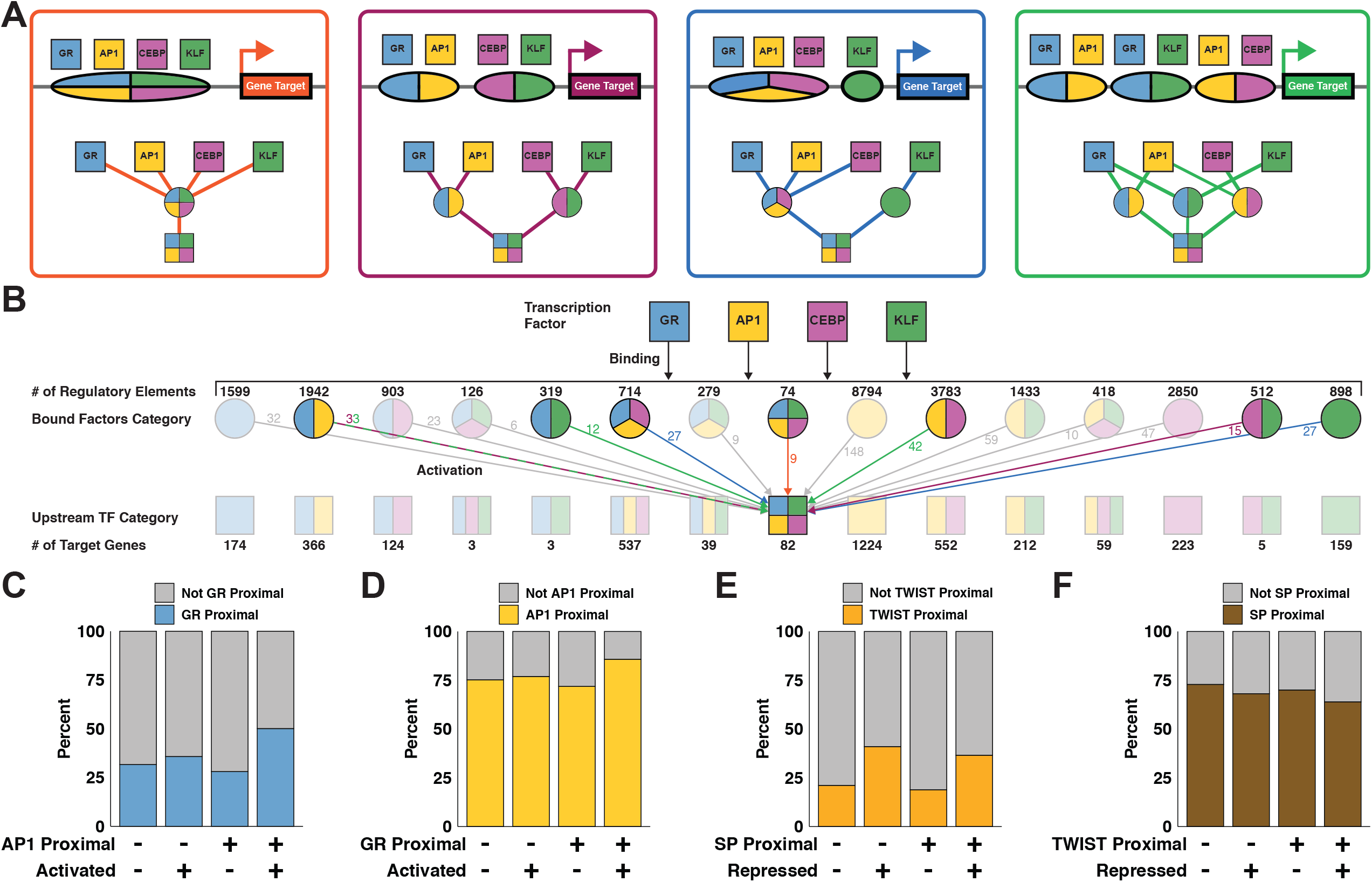
TFs bind REs and activate genes individually and combinatorially. A) Schematics and modular networks illustrate 4 variations in AP1, CEBP, GR, and KLF binding patterns that can lead to activation of gene targets: all 4 factors bind to a single RE (orange), 2 REs each bound by 2 factors (purple), 3 factors bind one RE and 1 factor binds another (blue), and a more complex combination with redundant factor contributions at multiple REs (green). B) A wide network depicts cooperation between TFs. Top squares indicate GR, AP1, CEBP, and KLF TF families. Second row circles represent 15 classes of REs each bound by a different combinations of factors. Third row squares represent 15 classes of genes each regulated by a different combination of factors. There are multiple potential combinations of REs that can produce the same gene class, as illustrated in (A). The middle square representing the 82 genes activated by all 4 factors is an example of variable regulatory combinations. Colored arrows and numbers correspond to RE combinations as depicted in (A). C) Genes were sorted into categories based on proximity to AP1 motifs, proximity to GR motifs, and activation status. We calculated 95% confidence intervals for odds ratios based on contingency tables consisting of genes stratified by their proximity to inferred AP1 and GR regulatory elements and by whether the genes were dynamic. The confidence interval for genes that are not proximal to inferred AP1 regulatory elements is 0.97 to 1.49. The confidence interval for genes that are proximal to inferred AP1 regulatory elements is 2.3 to 2.89. The increase in odds ratio prediction suggests that activated genes proximal to AP1 are significantly more likely to be proximal to GR than non activated genes. D) The confidence interval for genes not proximal to GR is 0.96 to 1.27. The confidence interval for genes proximal to GR is 1.94 to 2.87. Again, the increase indicates that AP1 and GR factors coordinate to activate transcription. E) The fraction of repressed genes proximal to TWIST increases regardless of the presence of SP with odds ratio confidence intervals of 2.2-3.07 and 2.21-2.79 when not proximal and proximal to SP peaks respectively. F) We find a lower proportion of repressed genes proximal to SP motifs both in the presence and absence of TWIST. The odds ratio confidence intervals do not change with the presence of TWIST, going from 0.71-0.89 to 0.64-0.9 when in proximity to predicted TWIST REs.

Interestingly, there is not a significant relationship between magnitude of RE accessibility change and number of regulatory factors (Figure S6C). We found that the relative change in transcription positively correlates with the number of immediate upstream activators in the network (Figure S6D). Normalizing transcriptional change by local accessibility change eliminates the observed correlation between transcription and number of regulatory factors (Figure S6E). We confirmed this observation by plotting transcription of all predicted target genes against total local accessibility stratified by the number of regulatory factors and peaks (Figure S7). We find that more local regulatory peaks, which corresponds greater total local accessibility, correlates with a greater magnitude of transcription. However, the number of regulatory factors largely does not affect transcription. Therefore, we find that transcription is positively correlated with total local accessibility change, regardless of the number of factors effecting that change. We conclude that if a gene is regulated in the network, the magnitude of expression change is independent from the number of upstream TFs.

Since the degree of activation and repression are unrelated to the number of upstream factors, we asked if having multiple TFs upstream in the network influences whether a gene is dynamic. To determine whether two TFs cooperate with one another we considered genes close to dynamic peaks with either a single TF motif or both TF motifs. We determine if the fraction of dynamic and nondynamic genes proximal to a TF is influenced by the presence of another TF. We define TF-proximal genes as genes that are close to dynamic ATAC peaks containing the TF motif. We find that there is no difference between the fraction of GR-proximal activated genes in the absence of AP1. However, there is an increase in the fraction of activated genes proximal to GR in the presence of AP1 (Figure 6C). The reciprocal analysis shows that AP1 is a more effective activator in the presence of GR (Figure 6D). These results support the model that AP1 and GR coordinate with one another to increase the likelihood of gene activation. The repressive factors TWIST and SP do not seem to work together in this way. The fraction of repressed genes proximal to TWIST increases regardless of the presence of SP (Figure 6E). This suggests that TWIST functions largely independently of SP, supporting our hypothesis that the two TFs result in gene repression through unrelated mechanisms (Figure S3B & C). Interestingly, we find a lower proportion of repressed genes proximal to SP motifs both in the presence and absence of TWIST (Figure 6F). We speculate that these genes tolerate dissociation of SP and maintain their expression levels despite local decreases in chromatin accessibility. In support of this explanation, we find higher basal transcription and lower magnitude of repression in genes proximal to SP, suggesting these genes are more actively transcribed before loss of SP (Figure S6F & G). These results highlight the complexity of gene regulatory control and how kinetic networks reveal coordinate and independent relationships between transcription factors.

### Multi-wave networks incorporate molecular dynamics and kinetic information

We further interrogate the adipogenesis gene regulatory network by leveraging temporal information to infer multiple waves of accessibility and transcriptional changes throughout the time course. The importance of TFs can be inferred by the number of predicted direct target genes (Figure 6) or the total number of connected downstream genes. The latter is captured by temporal multi-wave network depictions. We assembled a representative multi-wave deep network (Figure 7A). The differentiation cocktail induces AP1 and GR binding to thousands of REs to activate thousands of genes; binding at 4 of these REs results in activation of the *Twist2* gene (Figure 7A & B). The resulting TWIST2 protein returns to the nucleus and binds hundreds of REs and represses its target genes. Among the hundreds of repressed TWIST2 target genes are the late (40+ minutes) acting factors *Sp1* and *Sp3* (Figure 7C). The decreased occupancy of SP transcription factors from the genome leads to decreases in RE accessibility and attenuation of gene expression. Our network suggests that SP1/3 dissociation and TWIST2 binding leads to repression of *Srf* (Figure 7D). We hypothesize that if we were to extend the time course, we would identify the SRF binding motif in REs decreasing in accessibility beyond 4 hours as result of attenuated transcription.

**Fig. 7.**
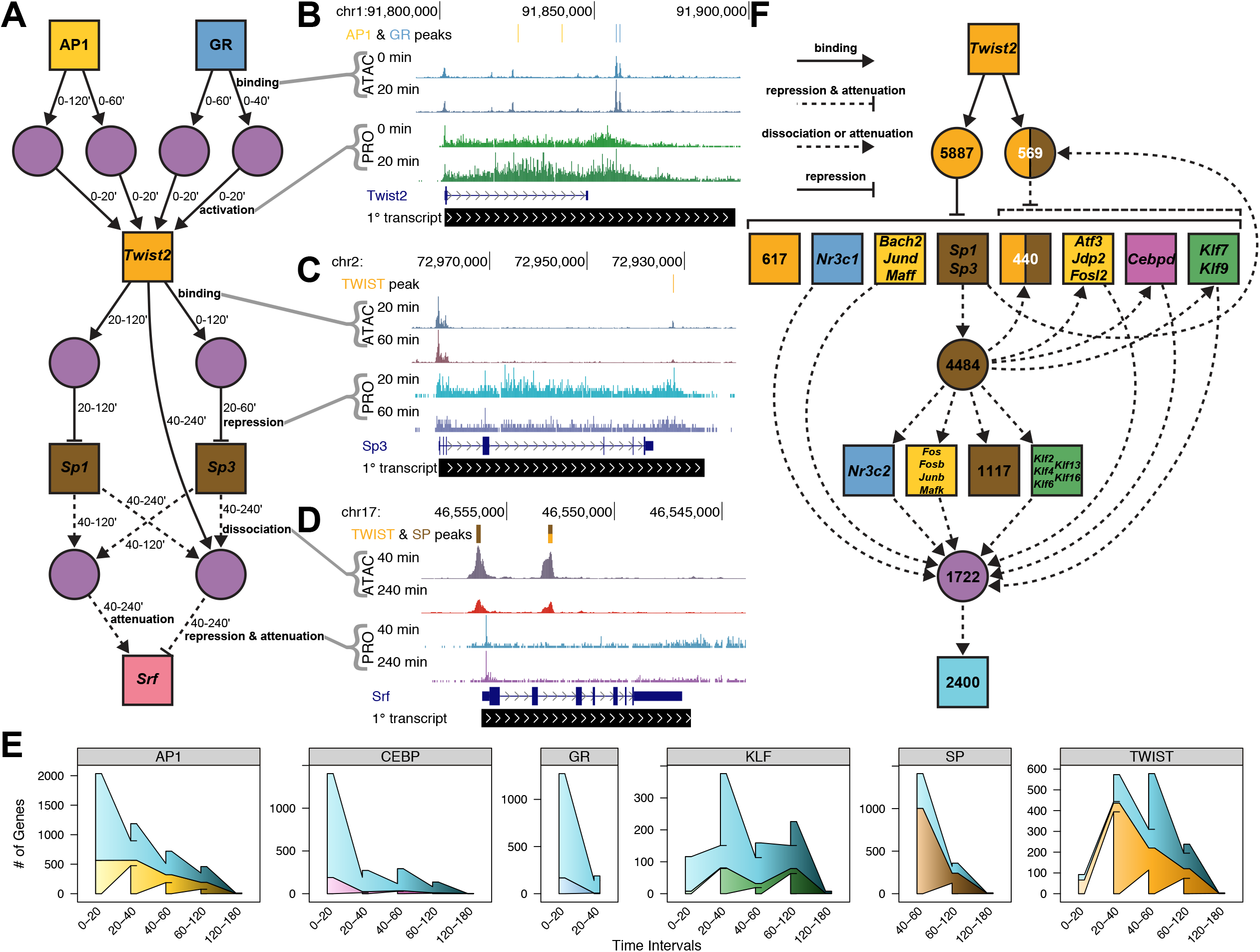
Variations in TF gene expression lead to downstream changes in accessibility and transcription. A) A deep network highlights temporal components of the regulatory cascade. Node and edge characteristics are illustrated as in Figure 4. In addition, we added time interval attributes to respective edges to indicate when REs and genes are changing. B-D) UCSC genome browser shots indicate accessibility and peaks (top three tracks) as well as nascent transcription (next two tracks) dynamics between the indicated time points. E) Wedged bar plots quantify the regulatory kinetics across the time course for indicated factors. The x-axis intervals represent the time range in which the indicated number (y-axis) of genes are regulated (connected in the network) by the specified factor. Wedges between bars indicate carryover elements from previous time interval and the outer “wings” represent elements that are not included in the previous time interval. The top shaded blue wedges represent genes regulated by multiple factors; bottom wedges represent genes that are solely regulated by the indicated factor. F) A *Twist2*-centric network illustrates the high connectivity and influence of *Twist2*.

Many genes activated in the early phase of the time course are repressed later on, either through active repression or factor dissociation. We detect these negative feedback loops for each activating TF (Figure S8A). About 63% of AP1, 74% of CEBP, and 80% of GR *cis*-edges are transient. Only 27% of KLF *cis*-edges are attenuated, suggesting that KLF-mediated activation is less transient and less dependent on the extracellular stimuli found in the adipogenic cocktail. Similarly, a minority of TWIST *cis*-edges and no SP *cis*-edges are attenuated, indicating that SP and TWIST factors mediate sustained repression. A much smaller proportion of *trans*-edges are attenuated, implying that accessibility changes downstream of factor binding and dissociation are more stable (Figure S8B) than changes in nascent transcription.

We find that regulatory potential for each TF varies greatly throughout the time course. AP1, CEBP, and GR activate the most genes during the initial phase of the time course, indicating that these TFs precipitate the initial wave of signaling during the first 20 minutes. Transcriptional activation of TWIST and KLF family genes by the initial factors leads to the next wave of signaling after 20 minutes (Figure 7B and Figure S8C). We begin to detect changes in accessibility at KLF- and TWIST-bound REs as early as 20 minutes (Figure S8D), however these presumptive binding events do not manifest as detectable changes in nascent transcription until 40 minutes (Figure 7E). Although we had originally expected changes in accessibility and transcription to be observed concomitantly, these data show that we have the sensitivity to detect changes in RE accessibility before changes in transcription.

In addition to the TFs whose activity is stimulated by the adipogenesis cocktail, we identify transcriptionally regulated TF genes that are highly connected nodes within the network. The *Twist2* gene is the most highly connected node and directly affects accessibility and transcription of thousands of downstream nodes by binding REs and repressing proximal genes (Figure 7F). TWIST2 acts through intermediate factors, such as SP, AP1, GR, to repress thousands of additional genes. In the case of SP, TWIST2 mediated repression of *Sp1* and *Sp3*, results in SP dissociation and activation attenuation of downstream genes. TWIST2-mediated repression of AP1 factors causes AP1 dissociation and attenuation of AP1-mediated activation. The cumulative result from both direct TWIST2 action and indirect dissociation / attenuation of TWIST2-targeted TF families affects accessibility at 12662 REs and 4574 genes. We believe that TWIST2 may have been overlooked as an important adipogenic TF because *Twist2* is only transiently activated (Figure S3C), but this kinetic network implicates TWIST2 as a critical intermediary in the adipogenesis cascade.

### TWIST2 represses predicted target genes

We tested whether inferred TWIST2-repressed target genes from the network increase expression upon *Twist2* depletion. We used two different shRNA sequences (v1 & v2) to knockdown *Twist2* and harvested RNA for RNA-seq at 0, 1, 2, and 4 hours after switching cells into differentiation media. We observed a *∼* 50%-75% reduction of *Twist2* expression prior to differentiation (Figure 8A & Figure S9A).Unlike PRO-seq, RNA-seq requires mature mRNA accumulation above baseline signal to detect activation and RNA degradation to detect repression. Therefore many observed transcriptional changes at nascent RNA level take much longer to be detected at the mature RNA level. For RNA-seq analysis we only focused on the 520 predicted TWIST2 targets that are significantly repressed in both the PRO-seq and RNA-seq time courses. The other 547 predicted TWIST2 target genes from the network are not detected as repressed by conventional RNA-seq, so we would have no power to detect derepression upon *Twist2* depletion. Approximately 66% of the examined genes were expressed at higher levels at baseline in the *Twist2* knockdown as compared to the control, supporting our hypothesis that TWIST2 directly represses the majority of our predicted targets (Figure 8B& Figure S9B). This result is likely an underestimate of the specificity of our network, because the cells can compensate for chronic RNAi-mediated depletion of *Twist2*.

We then measured the effect of chronic *Twist2* depletion on differentiation-induced transcription. *Twist2* was activated in both the control and the knockdown samples, supporting our PRO-seq results (Figure 8A & Figure S3C). Interestingly, *Twist2* was more strongly activated in the knockdown sample than in the control (*∼*3.1-fold to *∼*2-fold).This is likely because the RNAi machinery cannot keep up with the dynamic accumulation of *Twist2* transcripts in the hours following treatment with the differentiation cocktail. Since we inferred that TWIST2 represses target genes in this system, then we would expect a greater degree of repression over the time course in the knockdown samples due to the relatively greater accumulation of TWIST2 protein. This analysis more directly tests the accuracy of predicting TWIST2-target genes compared to chronic knockdown. We find that *∼*75% of the predicted targets are repressed to a greater magnitude in the shTwist2 samples compared to the control knockdown (Figure 8C). We observe supporting results with a second, less effective knockdown (Figure S9C). The above findings support the conclusions from the network and indicate that TWIST2 is a transcriptional repressor of predicted target genes in 3T3-L1 differentiation.

**Fig. 8.**
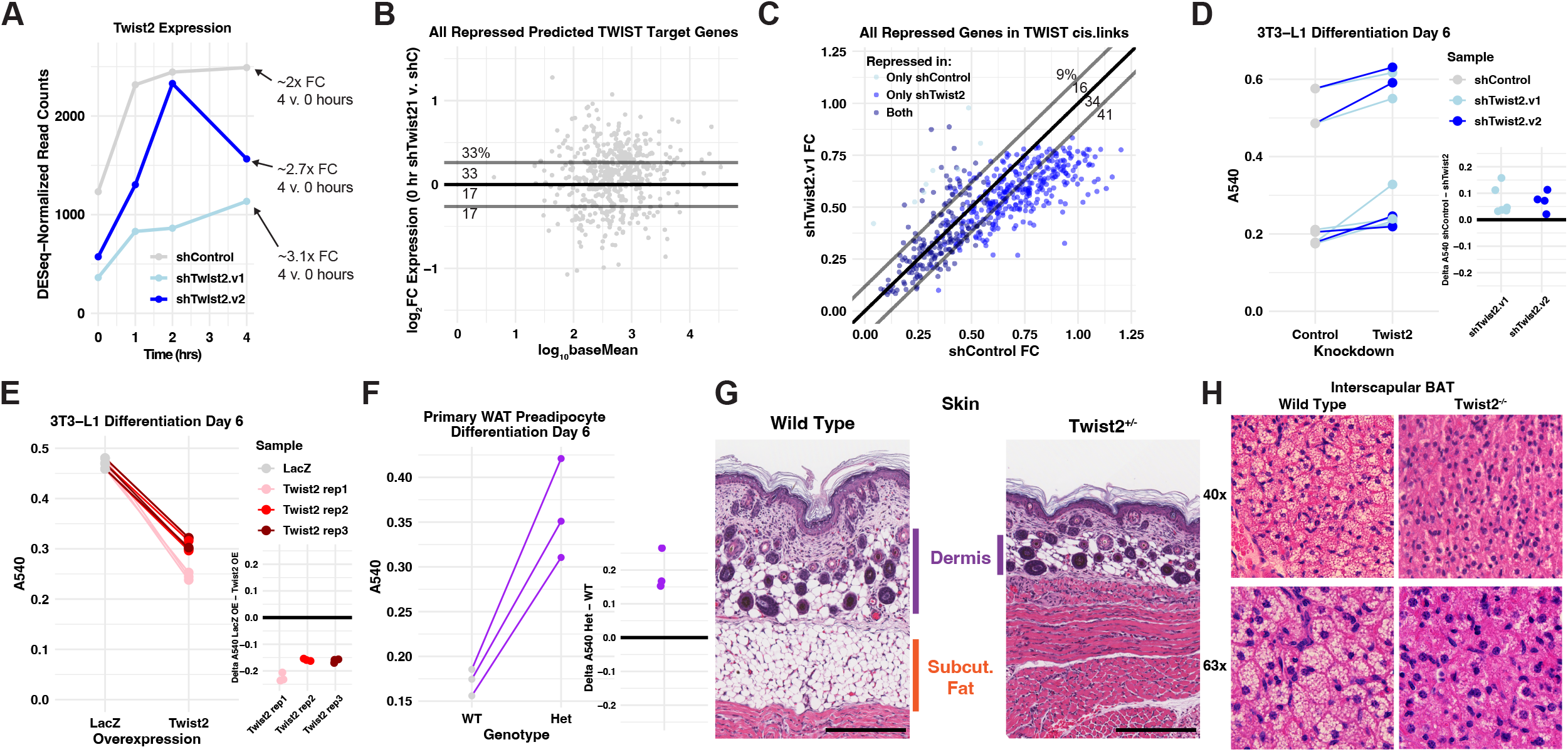
TWIST2 represses target genes and inhibits adipogenesis. A) *Twist2* expression as measured by RNA-seq at indicated time points following differentiation initiation. shTwist2.v1 is a single shRNA construct while shTwist2.v2 is a combination of shRNAs targeting *Twist2*. B) Expression of predicted TWIST2 target genes that are repressed in the RNA-seq time course before addition of the adipogenic cocktail. Y-values indicate fold change of gene expression in *Twist2* knockdown versus control. X-values indicate basal expression of each gene. The majority of genes are expressed to a higher degree in the knockdown samples, suggesting that loss of basal TWIST2 leads to derepression of target genes. C) Each data point represents one of the 520 predicted TWIST2 target genes that are repressed in the RNA-seq time course. We identified the time point comparison for which each gene exhibits the greatest degree of repression. X-values represent fold change for the relevant comparison in the shTwist2 samples and Y-values represent fold change for the same comparison in the shControl samples. Predicted TWIST2 target genes largely exhibit a greater degree of repression in the shTwist2 samples due to a relatively greater magnitude of *Twist2* activation. D) shControl or shTwist2 3T3-L1s were stained with Oil Red O after 6 days of differentiation. Absorbance at 540nm represents total lipid uptake of the cell population. Lines connect control and experimental samples from the same experimental replicate. Inset plot displays the difference in absorbance between the indicated shRNA treatment and the control knockdown for each replicate. TWIST2 depletion causes an increase in fat uptake, implicating the TF as a negative regulator of differentiation. E) 3T3-L1s overexpressing either LacZ or TWIST2 were stained with Oil Red O after 6 days of differentiation. TWIST2 overexpression causes a decrease in fat uptake, supporting the conclusion that TWIST2 is a negative regulator of adipogenesis. Lines and inset plot are used as in (D). F) Primary WAT preadipocytes harvested from P3 mice were induced to differentiate for 6 days and stained with Oil Red O. We pooled preadipocytes from animals with the same genotype. Wild type and heterozygous samples consist of preadipocytes pooled from 2 and 6 animals respectively. The pooled samples were plated in triplicate for the experiment. Preadipocytes harvested from *Twist2*^*+/-*^ pups exhibit increased Oil Red O staining compared to those harvested from wild type mice. G) Hematoxylin and eosin staining of skin shows collapse of dermal (purple bars) and subcutaneous WAT (orange bar) in P3 *Twist2*^*+/-*^ mice compared to wild type. Scale bars indicate 200 µm. H) Hematoxylin and eosin staining of interscapular brown fat shows reduced fat droplets (large white/light colored circles) in P14 *Twist2*^*-/-*^ mice compared to wild type. Images taken at either 40x (top) or 63x (bottom) magnification.

### TWIST2 influences differentiation of 3T3-L1s and primary preadipocytes

To test TWIST2’s effect on differentiation of preadipocytes, we depleted TWIST2, induced differentiation of 3T3-L1s, and measured lipid uptake after 6 days of differentiation. We stained differentiated adipocytes with Oil Red O and measured absorbance at 540nm to quantify lipid uptake. Lipid accumulation is a cellular phenotype that acts as a proxy measurement for adipogenesis. Lipid uptake increased with shTwist2 treatment compared to shControl within each experiment, suggesting that TWIST2 expression negatively regulates differentiation in the 3T3-L1 system (Figure 8D). As an orthogonal approach, we designed and transduced a tetracycline-inducible 3xFLAG-tagged human *Twist2* construct into 3T3-L1 cells (Figure S9D). After 6 days of differentiation, 3T3-L1s overexpressing *Twist2* exhibited decreased Oil Red O staining (Figure 8E & Figure S9E), supporting our previous finding that TWIST2 expression reduces 3T3-L1 differentiation.

Next, we extracted preadipocytes from inguinal white adipose tissue (WAT) of 3 day old *Twist2*^*+/-*^ pups. We induced differentiation in the primary preadipocytes and found that preadipocytes derived from heterozygous mice differentiated to a greater extent than those derived from wild type mice (Figure 8F & Figure S9F). The 3T3-L1 and primary cultured preadipocyte results indicate that TWIST2 opposes induced differentiation in both *in vitro* and *ex vivo* contexts.

We found that *Twist2*^*+/-*^ mice have a deficiency of dermal and subcutaneous white adipose tissue in the skin (Figure 8G). *Twist2*^*-/-*^ mice have fewer and smaller fat droplets within interscapular brown adipose tissue (BAT) deposits (Figure 8H). Other groups have reported loss of subcutaneous fat and a paucity of fat storage in *Twist2*^*-/-*^ mice (Kim et al. 2022; Šošić et al. 2003; Tukel et al. 2010). We postulate that TWIST2 acts as a ‘brake’ on adipogenesis, preventing cell exhaustion and apoptosis during the differentiation process. Regulated braking of adipogenesis may be necessary to allow supportive adipose tissues to sufficiently develop in the mouse. Isolated preadipocytes may be able to overcome the additional stress *in vitro*, but not within their native tissue context.

## Discussion

Kinetic accessibility and nascent transcriptional profiling of developmental cascades can identify key regulatory nodes that may be transiently active, but are nonetheless necessary for proper cellular differentiation. We present an extremely rapid and precise capture of chromatin and transcription changes induced by an adipogenic cocktail. These changes represent the first few waves of differentiation signaling and precipitate the cellular transition process. RE accessibility and gene transcription change within minutes of initiating adipogenesis. By focusing only on dynamically accessible REs, we can infer TF binding and dissociation events that drive adipogenesis without performing hundreds of genomic ChIP experiments. We find a multitude of enriched TF family motifs, many of which have been previously associated with adipogenic REs including AP1, GR, KLF, and CEBP (Siersbæk et al. 2014). We do not identify PPAR*γ*, the master regulator of adipogenesis (Lefterova et al. 2014; Rosen et al. 2002), as a driver of adipogenic signaling. This agrees with previous conclusions that PPAR*γ* does not influence adipogenesis until several days into the process (Nielsen et al. 2008). Stable PPAR*γ* activity is indispensable for adipogenesis and maintaining adipocyte identity, but other factors may be critically important and overlooked because their role is transient.

Our method implicates TWIST2 as a novel contributor to adipogenesis. The TWIST subfamily of bHLH TFs homo- and heterodimerize with other bHLH proteins to affect gene expression. Although TWIST family factors all recognize the same DNA motif, different members can act as either activators or repressors. TWIST proteins can repress transcription by non-productive dimerization with TWIST family activators, competing with TWIST family activators for DNA motifs, or by recruiting chromatin condensers like HDACs to the genome (Bialek et al. 2004; Gong and Li 2002; Hamamori et al. 1999; Hayashi et al. 2007; Koh et al. 2009; Lee et al. 2003; Šošić et al. 2003). Previous studies have implicated TWIST2’s role in targeting corepressors (Fu et al. 2011; Kim et al. 2022). While multiple mechanisms may be at play in our system, we hypothesize that our observed repressive effects are downstream of increased TWIST2 binding. TWIST1 and TWIST2 negatively regulate multiple developmental pathways including myogenesis, osteogenesis, and myeloid differentiation (Bialek et al. 2004; Gong and Li 2002; Hebrok et al. 1994; Murray et al. 1992; Sharabi et al. 2008; Spicer et al. 1996). The role of the TWIST TF family in adipogenesis is less clear. While TWIST1 and TWIST2 are known regulators of mature adipose tissue homeostasis, TWIST1 does not affect adipogenesis (Dobrian 2012; Lee et al. 2003; Pan et al. 2009). Interestingly, homozygous *Twist2* mutations cause Setleis syndrome, a disease characterized by facial lesions lacking subcutaneous fat (Tukel et al. 2010). *Twist2* knockout mice develop such lesions and lack lipid droplets within the liver and brown fat tissue (Figure 8G & H) (Šošić et al. 2003; Tukel et al. 2010). Our *in vitro* and *ex vivo* data indicate that TWIST2 acts as a negative regulator of adipogenesis. TWIST2’s immediate activation in 3T3-L1 differentiation therefore indicates a negative feedback mechanism to slow differentiation. The TWIST family is a key regulator of the epithelial-mesenchymal transition, further supporting our observation that TWIST2 prevents 3T3-L1 differentiation **??**. The loss of this negative feedback may result in cell death, leading to the absence of adipose tissue observed *in vivo*. Even with a well-studied system such as adipogenesis, these methods were able to identify *Twist2* as a novel regulator of the differentiation cascade.

Our networks define genes sets that are predominantly regulated by a single TF. We can track changes in RNA polymerse density within the gene sets to identify the target regulatory steps of individual TFs. Stimulated pause release is an established cause of early gene activation in adipogenesis (Wang et al. 2021). We find that GR is largely responsible for the observed increase in pause release. GR is a well-established activator of gene expression (Vockley et al. 2016), often in combination with AP1 (Biddie et al. 2011). Other activating factors, including AP1, increase RNA polymerase recruitment to the gene. By acting on separate steps, GR and AP1 provide non-redundant stimuli to target genes. We find GR and AP1 are conditionally dependent upon one another in their potential to activate local genes. A recent study suggests that both AP1 and CEBP act as pioneer factors that prime the genome for GR-induced transcription activation (Wissink et al. 2021). We find all three of these factor families activate the initial wave of transcription changes, both in combination and in isolation.

We confidently differentiate primary, secondary, and tertiary transcriptional changes by examining multiple, closely spaced time points upon induced adipogenesis. ATAC-seq, ChIP-seq, or chromatin conformation assays alone can only suggest functional relationships between REs and genes (Lieberman-Aiden et al. 2009; Ren et al. 2000). Similarly PRO-seq and RNA-seq return transcription changes with little information regarding upstream regulation. We define individual regulatory relationships more directly by focusing only on the ATAC peaks and genes that are proximal and covary in their dynamics (i.e. significantly change over in the same direction the time course). Our bipartite directed graph networks are unique in the gene regulation field because each edge represents a functional interaction as opposed to an abstract relationship between linked nodes. *Trans* edges represent binding of TF proteins to cognate DNA elements and *cis* edges describe regulatory interactions between REs and target genes. These networks can define genes sets that are predominately regulated by a single TF and identify the target regulatory steps of the TF. Highly connected nodes in the network are candidate key regulatory hubs in the differentiation cascade. Moreover, these networks ascribe time attributes to each edge, so subgraphs that respect the flow of time are easily extracted from the larger graph. This integrative genomics approach to network construction can be applied to a multitude of cellular responses and transitions to uncover novel biology and new hypotheses.

## Methods

### 3T3-L1 culture and differentiation

3T3-L1 cells were provided by Thurl Harris. 3T3-L1 cells were cultured in high glucose DMEM (Gibco) supplemented with 10% newborn calf serum, 1% fetal bovine serum (FBS), 100 U/mL Penicillin G, and 100 *µ*/mL streptomycin. We induced adipogenesis *∼* 3 days after cells reached confluency by switching cells into high glucose DMEM supplemented with 0.25 *µ*M Dexamethasone, 0.5 mM 3-isobutyl-1-methylxanthine, 2.5 units/mL insulin, 10% FBS, 100 U/mL Penicillin G, and 100 *µ*g/mL streptomycin (Bernlohr et al. 1984; Green and Kehinde 1974). We collected enough cells at the indicated time points for three replicates of ATAC-seq and PRO-seq.

### shRNA-Mediated Knockdown

We purchased lentiviral shRNA-expressing constructs targeting Twist2 (Millipore Sigma clone IDs TRCN0000086084, TRCN0000086085, TRCN0000086086) and a non-mammalian control (Millipore Sigma SHC202). HEK-293T cells were transfected with shRNA constructs and lentiviral packaging constructs pMD2.G (Addgene #12259) and psPAX2 (Addgene #12260). We isolated and filtered supernatant after 24 and 48 hours. We transduced 3T3-L1s with virus in 8 µg/mL polybrene. Cells were switched into puromycin-containing selection media after 48 hours. After another 48 hours surviving cells were plated for further experiments.

### Oil Red O staining

For Oil Red O assays 3T3-L1s were cultured and differentiated as described above. During differentiation media was changed every 2 days. At day 6 of differentiation we stained cells with Oil Red O as previously described (Kraus et al. 2016). Briefly, cells were washed with PBS, fixed with 4% formaldehyde for 15 minutes, and stained with a 0.2% Oil Red O and 40% 2-propanol solution. After incubating for 30 minutes adipocytes were washed five times with distilled water and the dye was eluted with 2-propanol. Absorbance was measured at 540 nm using 2-propanol as a blank.

### RNA extraction and RNA-seq

3T3-L1s were cultured and differentiated as described above. At the indicated time points we harvested cells and extracted RNA using the Direct-zol-96 RNA kit (Zymo #11-331H). Samples were then sent to Novogene for bulk RNA-seq.

### Immunoblotting

3T3-L1s were cultured and differentiated as described above. Immunoblotting was performed on 7.5 µg RIPA lysate as previously described (Janes 2015). Samples were electrophoresed through 1.5-mm thick 12% polyacrylamide in tris-glycine running buffer (25 mM tris base, 250 mM glycine, and 0.1% SDS) at 130 V for 90 minutes. Proteins were transferred to a PVDF membrane (Millipore; Immobilon-FL, 0.45 mm pore size) in transfer buffer (25 mM tris, 192 mM glycine, 0.0375% SDS, and 20% methanol) at 100 V for 1 hour on ice. Membranes were blocked with 0.5X Odyssey blocking buffer in TBS. Primary antibodies were diluted in 0.5X Odyssey blocking buffer + 0.1% Tween-20. The following primary antibodies were used: Flag (Millipore Sigma F1804, 1:2000), HSP90 (Santa Cruz Biotechnology sc-13119, 1:2000), Tubulin (Cell Signaling Technology 2148S 1:2000). Membranes were washed with TBS-T and exposed to fluorophore-conjugated secondary antibody diluted in 0.5X Odyssey blocking buffer+ 0.1% Tween-20 + 0.01% SDS. Following another round of washing, membranes were scanned on an Odyssey infrared scanner (LI-COR).

### Genome browser visualization

Genome browser (Kent et al. 2002) images were taken from the following track hub: http://guertinlab.cam.uchc.edu/adipo_hub/hub.txt. An identical track is reproduced on a public server: https://data.cyverse.org/dav-anon/iplant/home/guertin/adipo_hub/hub.txt

### ATAC-seq library preparation

We prepared ATAC-seq libraries as previously described (Corces et al. 2017). We trypsinized and collected cells in serum-free growth media. We counted *∼* 5 × 10^4^ cells perreplicate and transferred them to 1.5 mL tubes. We centrifuged cells at 500 x *g* for 5 minutes at 4°C and resuspended the pellet in 50 *µ*L cold lysis buffer (10mM TrisHCl, 10mM NaCl, 3mM MgCl_2_, 0.1% NP-40, 0.1% Tween-20, 0.01% Digitonin, adjusted to pH 7.4) and incubated on ice for 3 minutes. We washed the samples with 1 mL cold wash buffer (10mM Tris-HCl, 10mM NaCl, 3mM MgCl_2_, 0.1% Tween-20). We centrifuged at 500 x *g* for 10 minutes at 4°C and resuspended cells in the transposition reaction mix (25 *µ*L 2X TD buffer (Illumina), 2.5 *µ*L TDE1 Tn5 transposase (Illumina), 16.5 *µ*L PBS, 0.5 *µ*L 1% Digitonin, 0.5 *µ*L 10% Tween-20, 5 *µ*L nuclease-free water) and incubated at 37°C for 30 minutes. We extracted DNA with the MinElute kit (Qiagen). We attached sequencing adapters to the transposed DNA fragments using the Nextera XT Index Kit (Illumina) and amplified libraries with 6 cycles of PCR. We performed PEG-mediated size fractionation (Lis 1980) on our libraries by mixing SPRIselect beads (Beckman) with our sample at a 0.55:1 ratio, then placing the reaction vessels on a magnetic stand. We transferred the right side selected sample to a new reaction vessel and added more beads for a final ratio of 1.8:1. We eluted the final size-selected sample into nuclease-free water.

### ATAC-seq analyses

We aligned reads to the *mm10* mouse genome assembly with `bowtie2`, sorted output *BAM* files with `samtools`, and converted files to *bigWig* format with `seqOutBias` (Langmead and Salzberg 2012; Li et al. 2009; Martins et al. 2018). We called accessibility peaks with `MACS2` (Zhang et al. 2008). We sort reads into peaks using the `bigwig` R package and identify differentially accessible REs with `DESeq2` (Love et al. 2014; Martins 2014). We cluster dynamic peaks into response groups using `DEGreport` (Pantano 2019). We performed *de novo* motif extraction on dynamic REs with `MEME` (e-value cutoff of 0.01) and used `TOMTOM` (e-value cutoff of 0.05) to match motifs to the HOMER, Jaspar, and Uniprobe TF binding motif databases (Bailey et al. 2015; Heinz et al. 2010; Hume et al. 2015; Khan et al. 2018). We use the `bigWig` package to assess motif enrichment around ATAC-seq peak summits (Martins 2014).

### PRO-seq library preparation

We performed cell permeabilization as previously described (Mahat et al. 2016). We trypsinized and collected cells in 10 mL ice cold PBS followed by washing in 5 mL buffer W (10 mM Tris-HCl pH 7.5, 10 mM KCl, 150 mM sucrose, 5 mM MgCl_2_, 0.5 mM CaCl_2_, 0.5 mM DTT, 0.004 U/Ml SUPERaseIN RNase inhibitor (Invitrogen), protease inhibitors (cOmplete, Roche)). We permeabilized cells by incubating with buffer P (10 mM Tris-HCl pH 7.5, KCl 10 mM, 250 mM sucrose, 5 mM MgCl_2_, 1 mM EGTA, 0.05% Tween-20, 0.1% NP40, 0.5 mM DTT, 0.004 units/mL SUPERaseIN RNase inhibitor (Invitrogen), protease inhibitors (cOmplete, Roche)) for 3 minutes. We washed cells with 10 mL buffer W before transferring into 1.5 mL tubes using wide bore pipette tips. Finally, we resuspended cells in 500 *µ*L buffer F (50 mM Tris-HCl pH 8, 5 mM MgCl_2_, 0.1 mM EDTA, 50% Glycerol and 0.5 mM DTT). After counting nuclei, we separated cells into 50 *µ*L aliquots with *∼* 3-5 × 10^5^ cells each. We snap froze our aliquots in liquid nitrogen and stored them at -80°C. We prepared PRO-seq libraries as previously described (Sathyan et al. 2019). We included a random eight base unique molecular identifier (UMI) at the 5^*Õ*^ end of the adapter ligated to the 3^*Õ*^ end of the nascent RNA. We did not perform any size selection in an attempt to not bias our libraries against short nascent RNAs.

### PRO-seq analyses

First we used `cutadapt` to remove adapters from our reads (Martin 2011). We used `fqdedup` and the 3^*Õ*^ UMIs to deduplicate our libraries (Martins and Guertin 2018). Next we removed UMIs and converted reads to their reverse complement with the `FASTX-Toolkit` (Gordon 2010). As with the ATAC-seq samples, we used `bowtie2`, `samtools`, and `seqOutBias` to align, sort, and convert reads to *bigWig* files respectively (Langmead and Salzberg 2012; Li et al. 2009; Martins et al. 2018). We used `primaryTranscriptAnnotation` to adjust gene annotations based on our PRO-seq data (Anderson et al. 2020). We queried the *bigWig* files within the adjusted genomic coordinates with the `bigWig` R package and UCSC Genome Browser Utilities (Kent et al. 2010; Martins 2014). We identified differentially expressed genes with `DESeq2` (Love et al. 2014). We used `dREG` to define peaks of bidirectional transcription from our *bigWig* files (Wang et al. 2019). As with the ATAC-seq samples, we identified overrepresented motifs in dREG-defined REs with `MEME` and `TOMTOM` (Bailey et al. 2015). We evaluate motif enrichment around peak summits and polymerase density in the gene body and pause region with the `bigWig` package (Martins 2014). We define the summit of the pause peak for genes by first identifying the point of maximum density within 1 kb of the TSS. We define the pause region as the 50 bp window around the summit.

### DNA binding domain alignments

DNA binding domains were extracted from the TFClass database (Wingender et al. 2018) and TF paralogs that were absent from the database were extracted from the NCBI protein database. DNA binding domains were aligned using `FASTA` (Pearson and Lipman 1988) and the following command: `ssearch36 -s MD40 -m 8CBl`. Although there are six DNA motifs, the TWIST and ZNF families of DNA binding domains recognize the same motif, despite their lack of evolutionary conservation.

### Network construction

The bipartite directional networks with gene and RE nodes were inferred using a data-driven rules-based approach. The first rule to infer *trans*-edges from TF families to individual REs is that the RE must contain the cognate motif for the TF family. The second is that the peak must be dynamically accessible over some part of the time course. The third is that at least one gene encoding a member of the TF family must be expressed and activated (or in the case of SP, repressed) over the same time range. We restrict *trans*-edges attributed to GR to the first 40 minutes of the time course for reasons discussed in the text. Similarly, we do not draw edges from SP before 40 minutes. Next, we drew *cis*-edges between REs and proximal genes based on a different rule set. First, REs need to be within 10 kb of gene bodies as defined by primary transcript annotation of our PRO-seq data. We used `bedtools` to find gene-RE pairs that satisfied this rule (Quinlan and Hall 2010). Next, the peak and the gene need to covary in accessibility and transcription during the same time range. For example, a gene must be activated at the same time as its local RE is increasing in accessibility. We refined the distance requirements by incorporating constraints from our CDF analysis. For each activating factor (AP1, CEBP, GR, KLF) we find a set of pairwise comparisons within the time course for which factor REs are significantly closer to activated than nondynamic genes. We find a similar set of comparisons for repressive factors (SP, TWIST). For a gene to be linked to a factor RE with a *cis*-edge, we require that the gene must be dynamic in at least one of the comparisons identified by the CDF analysis for that factor. In addition, our CDF analysis also identifies the maximum distance between a factor RE and a regulated gene for each comparison. The RE and the gene’s TSS must be within the relevant distance threshold defined by the CDF.

### Compartment modeling

Detailed analysis and raw code is available at https://github.com/guertinlab/modeling_PRO_composites. We calculated pause region signal by summing the PRO-seq signal over a 50 base pair window centered on the summit of the pause peak. The gene body RNA polymerase density was determined by averaging the PRO-seq signal over the region from the end of the pause window to the transcription termination site determined by `primaryTranscriptAnnotation`. We described RNA polymerase density dynamics in each compartment using differential equations that incorporate rates of transcription initiation, premature termination, pause release, and elongation. We determined how different rate constants would affect hypothetical pause region and gene body densities and the pause index using the `pksensi` package.

We varied initiation, premature termination, and pause release rate constants from 0.01 to 1 molecules per second and varied elongation rate from 600-6000 bp/minute and calculated compartment density with each parameter set. We excluded any parameter estimates that result in more than 1 RNA polymerase molecule in the pause region at any given time. We selected all sets of parameters that resulted in the observed median composite pause index for all time points from Figure 4. Next, we sought to determine if changing the rate constants could explain observed changes in compartment density at different gene sets for the indicated time point comparisons. We varied rate constants from the early time point values to model RNA polymerase density ratio changes between time points. We varied the initiation and pause release constants from their initial values over a 5-fold range in each direction and determined which sets of constants produced the observed changes in pause density ratio for each gene. We allowed the target ratio to vary by 5% compared to the observed. We plotted model RNA polymerase density curves by choosing the set of parameters with the elongation rate closest to a consensus rate of 2500 bases / min (Ardehali and Lis 2009; Jonkers and Lis 2015), while still accurately reproducing the composite profiles densities within 5% of the original. In order to reproduce composite plots, we spread the pause density over a 50 base region and fit the density profiles to a waveform as previously described (Sathyan et al. 2019).

For the model, our only assumption is that elongation rate of a particular gene does not change between time points. In fact, if elongation rate was increasing in activated genes then we would expect a decrease in gene body PRO-seq signal. This finding has not been observed in other datasets. Of all the rate constants in the model, it is most reasonable to assume that the elongation rate stays constant while the others are affected by different regulatory mechanisms.

### Network construction

The bipartite directional networks with gene and RE nodes were inferred using a data-driven rules-based approach. The first rule to infer *trans*-edges from TF families to individual REs is that the RE must contain the cognate motif for the TF family. The second is that the peak must be dynamically accessible over some part of the time course. The third is that at least one gene encoding a member of the TF family must be expressed and activated (or in the case of SP, repressed) over the same time range. We restrict *trans*-edges attributed to GR to the first 40 minutes of the time course for reasons discussed in the text. Similarly, we do not draw edges from SP before 40 minutes. Next, we drew *cis*-edges between REs and proximal genes based on a different rule set. First, REs need to be within 10 kb of gene bodies as defined by primary transcript annotation of our PRO-seq data. We used `bedtools` to find gene-RE pairs that satisfied this rule (Quinlan and Hall 2010). Next, the peak and the gene need to covary in accessibility and transcription during the same time range. For example, a gene must be activated at the same time as its local RE is increasing in accessibility. In this context, REs / genes need to change accessibility / transcription by at least 10% over the time range to be considered activated or repressed. If the peak and gene are both activated or both repressed, then we draw a *cis*-edge between the RE and the gene. We refined the distance requirements by incorporating constraints from our CDF analysis. For each activating factor (AP1, CEBP, GR, KLF) we find a set of pairwise comparisons within the time course for which factor REs are significantly closer to activated than nondynamic genes. We find a similar set of comparisons for repressive factors (SP, TWIST). For a gene to be linked to a factor RE with a *cis*-edge, we require that the gene must be dynamic in at least one of the comparisons identified by the CDF analysis for that factor. In addition, our CDF analysis also identifies the maximum distance between a factor RE and a regulated gene for each comparison. The RE and the gene’s TSS must be within the relevant distance threshold defined by the CDF.

### Mouse experiments

All mouse experiments were performed in accordance with the relevant guidelines and regulations of the University of Virginia and approved by the University of Virginia Animal Care and Use Committee. Mice were housed in specific pathogen-free conditions under standard 12-h-light/dark cycle conditions in rooms equipped with control for temperature (21±1.5°C) and humidity (50±10%). Twist2 knockout mice were purchased from Jackson Laboratories (Strain 008712). Preadipocytes were isolated from inguinal WAT as previously described (Galmozzi et al. 2021) from P3 mice. Preadipocyte differentiation and staining was carried out as with 3T3-L1s. Interscapular skin and BAT samples were isolated from P3 or P14 mice respectively. Tissue samples were fixed with 4% formaldehyde. Fixed tissues were paraffin-embedded and H&E stained by the Research Histology Core at the University of Virginia.

## Data Access

All analysis details and code are available at https://github.com/guertinlab/adipogenesis. Raw sequencing files and processed *bigWig* files are available from GEO accession records GSE133147 (3T3-L1 PRO-seq), GSE150492 (3T3-L1 ATAC-seq), GSE219041 (C7 cells + dexamethasone PRO-seq), and GSE219051 (shTwist2 3T3-L1 RNA-seq).

## Competing Interest Statement

The authors have no competing interests to disclose.

## ACKNOWLEDGEMENTS

This work was funded by R35-GM128635 to MJG. T32-LM012416 supported ABD. We thank John Lukens for hosting animals. We thank Thurl Harris for providing 3T3-L1 cells and technical guidance. We thank Kevin Janes for providing reagents and suggestions. We thank Sathyan Mattada, Thomas Scott, Jacob Wolpe, Sailasree Rajalekshmi, and Theresa Gibney for critical feedback.

## AUTHOR CONTRIBUTIONS

ABD, BN, and MJG analyzed the data. NMW and FMD performed the 3T3-L1 ATAC-seq and PRO-seq experiments. ABD and DSL performed the 3T3-L1 and primary cell adipogenesis assays. RP, PP, and ABD performed the mouse tissue harvesting and preadipocyte extraction. LW designed and cloned the TWIST2 overexpression construct. ABD and MJG conceptualized and developed the project. ABD and MJG wrote the manuscript.

---

**Fig. S1.**
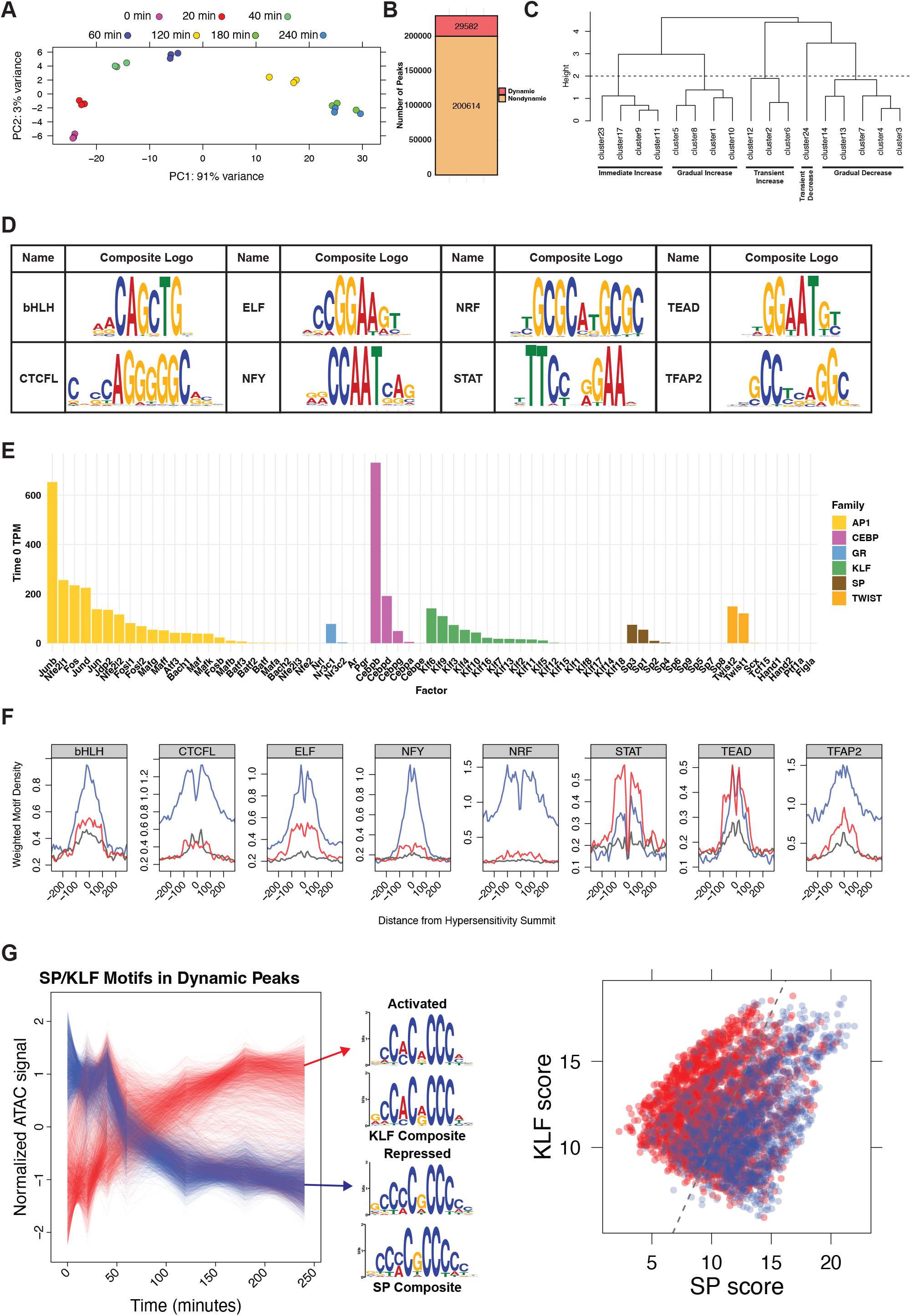
Detailed analyses of dynamic ATAC-seq peaks reveals potential regulatory transcription factors. A) Principal component analysis of ATAC-seq samples shows clear separation of samples by time. B) We used a likelihood ratio test to define ATAC-seq peaks as dynamic or nondynamic using an adjusted p-value cutoff of 1 * 10*™*8. C) After clustering ATAC-seq peaks based on accessibility dynamics, we combined clusters into more inclusive response classes by cutting the dendrogram at the indicated height of 2. D) In addition to the motifs discussed in Figure 1, *de novo* motif analysis identified 8 factor families as potential regulators of ATAC-seq peak dynamics. These motifs are less enriched than those in Figure 1. E) Expression of all members of potential regulatory TF families as measured by RNA-seq of 3T3-L1 cells before addition of differentiation media. F) We plotted enrichment of each motif family around peak summits as in Figure 1E. G) Composite motif extraction on all peaks with either SP or KLF motifs returns a KLF-like motif in the increased peaks and an SP-like motif in the decreased peaks, suggesting that KLF and SP factor families are associated with increased and decreased accessibility respectively. We scored each SP/KLF motif in a dynamic peak against an SP composite and a KLF composite and plotted the two scores against one another. Each data point is colored based on the dynamics of the peak. The dividing line separates points such that all motifs on the left are labeled as KLF motifs and all motifs on the right are labeled SP motifs. The line is drawn in such a way as to maximize the number of increased KLF peaks and decreased SP peaks.

**Fig. S2.**
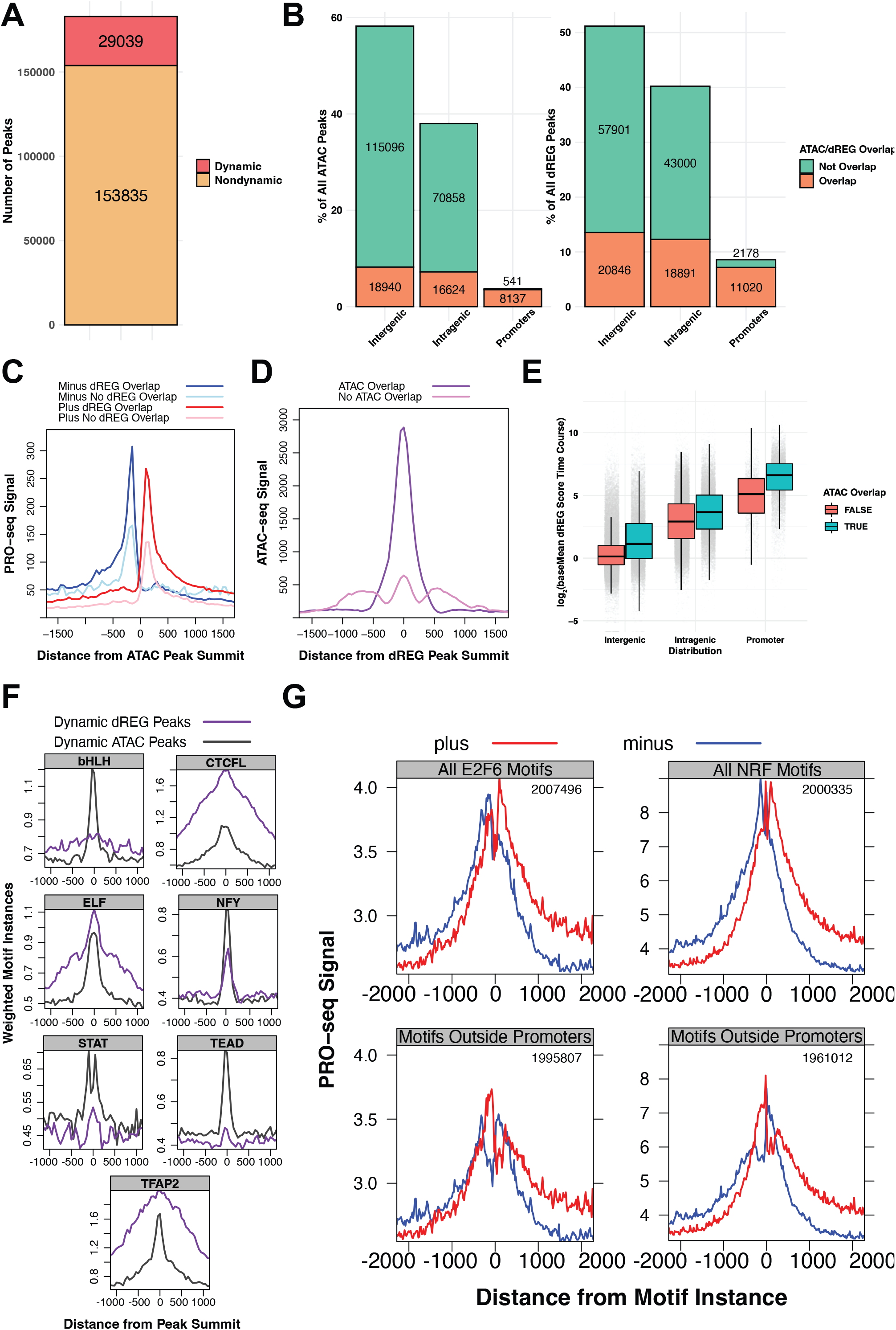
dREG peak analysis reveals potential factors driving bidirectional transcription. A) We used a likelihood ratio test to define dREG peaks as dynamic or nondynamic using an adjusted p-value cutoff of 1 * 10*™*5. B) ATAC-seq and dREG-defined REs - including dynamic and nondynamic peaks - largely overlap in promoter regions. Genome regions are defined as described in Figure 2 legend. Overlapping categories are not equal between ATAC and dREG peaks because multiple ATAC peaks may overlap a single dREG peak and vice versa. C) PRO-seq signal plotted around ATAC peaks separated by whether they overlap with dREG peaks. D) ATAC-seq signal plotted around dREG peaks separate by whether they overlap with ATAC peaks. E) dREG score for peaks sets stratified by location and ATAC overlap status. F) Motif density plots identify CTCF, ELF, and TFAP2 as enriched in dREG peak summits over ATAC-seq peak summits. Since these factors were not found in the *de novo* analysis, we did not consider them dREG-enriched factors. G) Plotting PRO-seq signal around non-SP dREG factor motifs shows the bidirectional transcription phenotype but not as clearly as with the SP motifs.

**Fig. S3.**
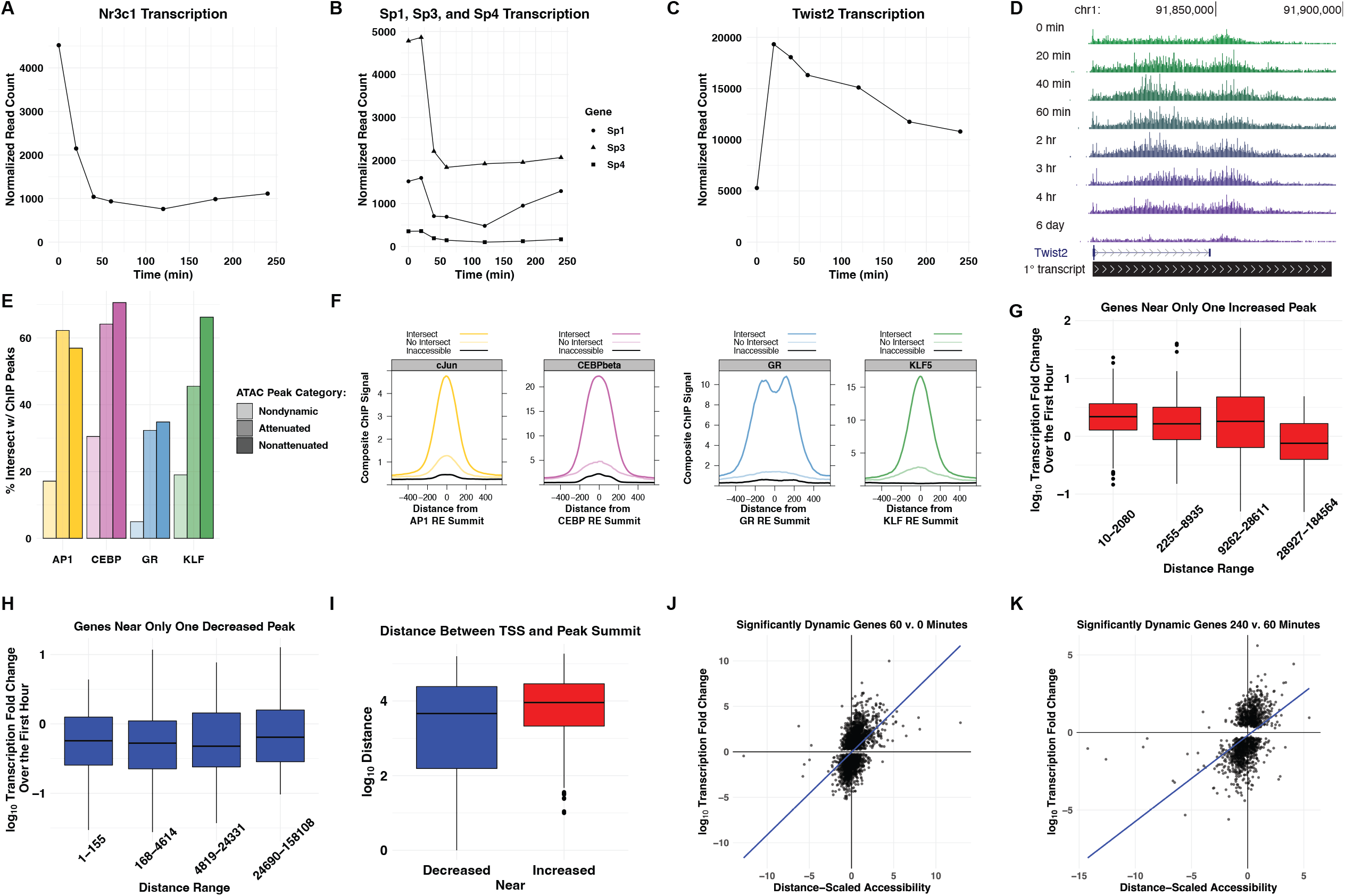
Gene transcription positively correlates with local accessibility dynamics. A-C) Normalized PRO-seq signal for A) the *Nr3c1* gene, B) the *Sp1, Sp3*, and *Sp4* genes, and C) the *Twist2* gene. *Nr3c1, Sp1, Sp3*, and *Sp4* all exhibit rapid early repression, while *Twist2* is immediately and transiently activated. D) The UCSC genome browser shot shows that early activation of *Twist2* gene has subsided by the time cells reach full maturity at 6 days. E) We compared our ATAC-seq derived REs with ChIP-seq peaks for AP1 (cJun and JunB), KLF (KLF4 and KLF5), CEBP*β*, and GR. ATAC-seq peaks are divided into nondynamic peaks that are not inferred binding events, attenuated peaks that increase early in the time course but decrease toward the end, and nonattenuated peaks that do not decrease in accessibility at any point. F) Composite ChIP-seq signal is plotted at indicated genomic regions. Regions are classified into dynamic REs that intersect with ChIP-seq peaks, dynamic REs that do not intersect with ChIP-seq peaks, and TF motifs that are not found in ATAC-seq peaks and are therefore inaccessible. G-H) Box and whisker plots representing genes within 10 kb of either (G) a single increased peak or (H) a single decreased peak. Genes were split into four quartiles based on the distance between the peak summit and the TSS of the gene. Box and whisker plots on the left contain the smallest distances, while those on the right represent the largest. The y-value is the transcription of the genes over the first hour of the time course. We find that for genes near increased peaks, the closer the peak is to the gene the more likely gene transcription and peak accessibility dynamics will covary. We do not find a similar relationship for genes near decreased peaks. I) Box and whisker plots representing genes within 10 kb of either a single increased peak (red plot) or a single decreased peak (blue plot). The y-value represents the distance between the peak summit and the TSS of the gene. Genes near decreased peaks are generally closer to the peak summit than the increased peak-gene pairs. J-K) Scatter plot of genes dynamically transcribed between J) 60 and 0 minutes or K) 240 and 60 minutes. Each data point represents a gene. The x-value is total accessibility change over the time period scaled by the distance between the TSS and each peak. Positive x-values indicate increases in local accessibility. The y-value is change in transcription over the same time period. A gene with positive x and y-values or negative x and y-values exhibits positive covariation between local accessibility and transcription dynamics. The majority of genes for both time point comparisons exhibit positive covariation, indicating a correlative relationship between accessibility and transcription. Chi-squared p-values are (J) 1.41 * 10^−98^ and (K) 4.35 * 10^−54^.

**Fig. S4.**
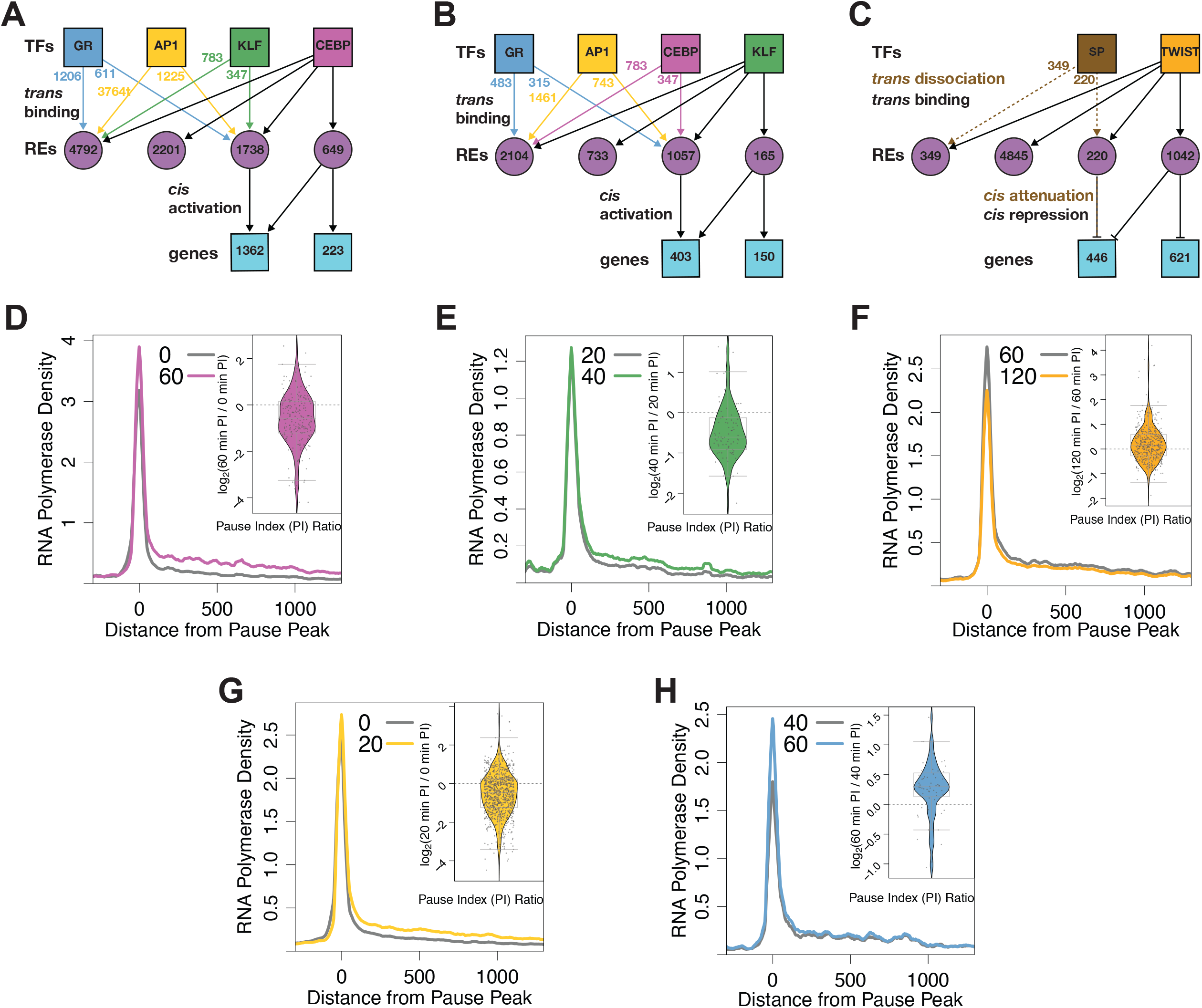
Modular networks downstream of CEBP, KLF, and TWIST identify genes and transcriptional steps regulated by individual factors. Modular networks downstream of (A) CEBP, (B) KLF, and (C) TWIST. Networks constructed as described in Figure 4. Composite PRO-seq signal plotted around pause peak summits of (D) 223 genes solely regulated by CEBP, (E) 150 genes solely regulated by KLF, (F) 621 genes solely regulated by TWIST, (G) 1224 genes solely regulated by AP1, and (H) 174 genes solely regulated by GR at the indicated time points illustrate average changes in RNA polymerse density within the pause region and gene bodies. Inset violin plots indicate pause indices as described in Figure 4.

**Fig. S5.**
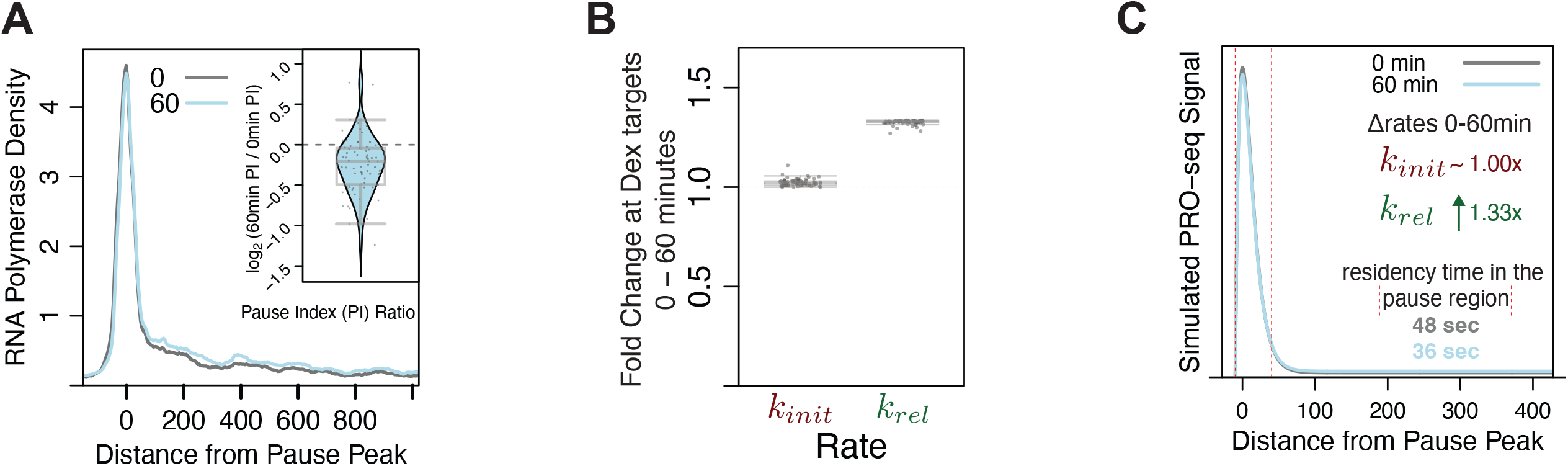
Dexamethsone-activated genes in C7 cells recapitulate findings at predicted GR target genes in 3T3-L1 differentiation. A) Composite polymerase density at 70 genes activated by dexamethasone treatment in C7 cells at indicated times. B) Compartment modeling of the activated genes holding *kpre* and *krel* constant. We find an approximate 1.33-fold increase in pause release rate explains the observed changes in polymerase density. C) A simulated composite derived from the parameters estimated in (B).

**Fig. S6.**
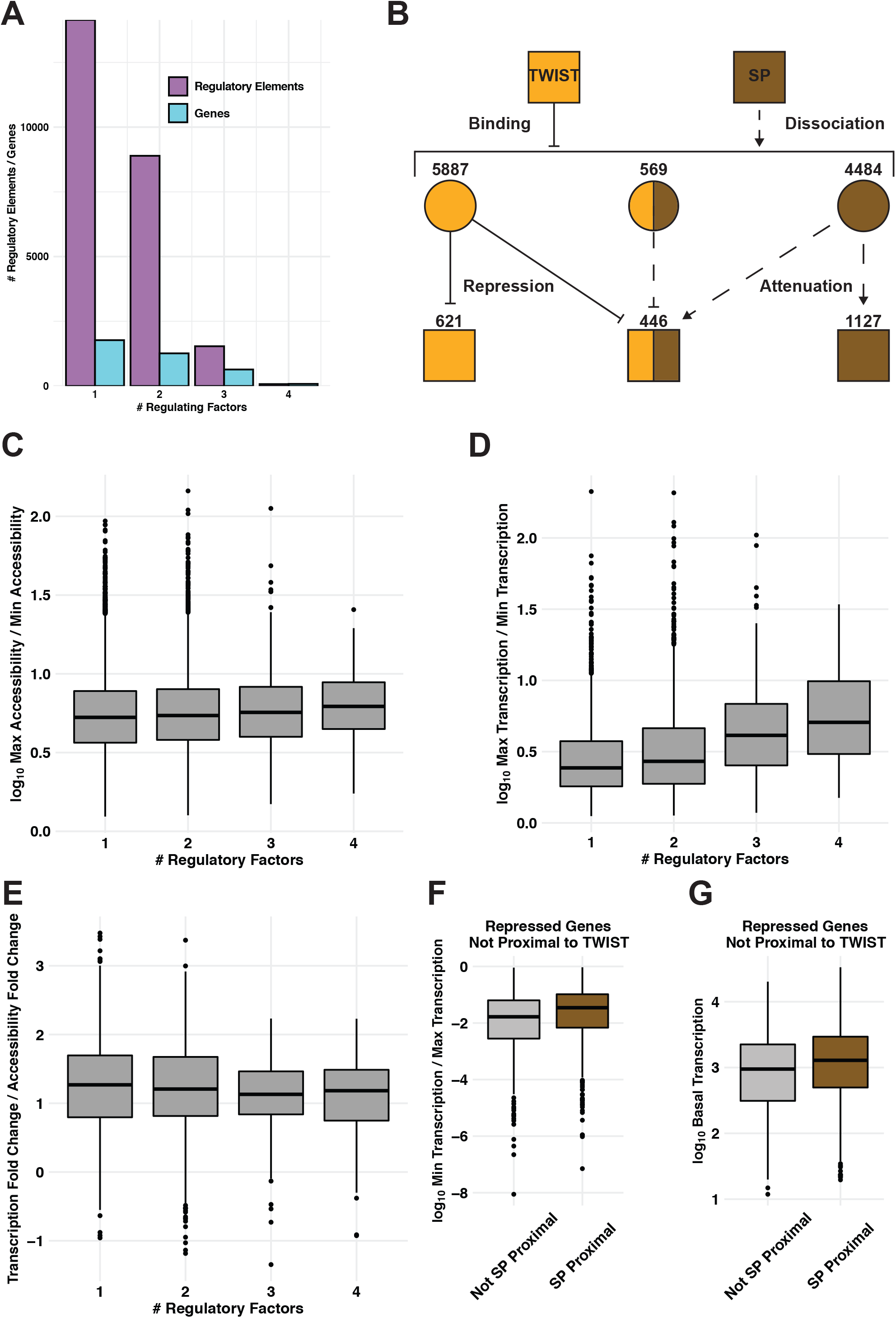
Total local accessibility influences magnitude of gene transcription changes. A) The bar plot quantifies the number of REs and genes that are activated by the indicated number of activating factors: GR, AP1, CEBP, or KLF. B) This network depicts REs and genes downstream of TWIST binding and repression and/or SP dissociation and attenuation. C) Data points represent all REs bound by AP1, CEBP, GR, and KLF. REs are stratified based on number of binding factors. Y-values are the fold change in accessibility. D-F) Data points represent all genes activated by AP1, CEBP, GR, and TWIST. D) Genes are stratified based on number of upstream regulatory factors. Y-values are the fold change in nascent transcription. There is a positive correlation between transcriptional change and number of regulatory factors. E) Genes are stratified as in (D), but y-values represent fold change in transcription divided by fold change in local accessibility. There is no correlation between normalized transcription and number of regulatory factors when controlling for changes in accessibility. F) Repressed genes proximal to SP motifs tend to exhibit a lower magnitude of transcriptional change. G) Repressed genes proximal to SP motifs tend to exhibit higher basal transcription.

**Fig. S7.**
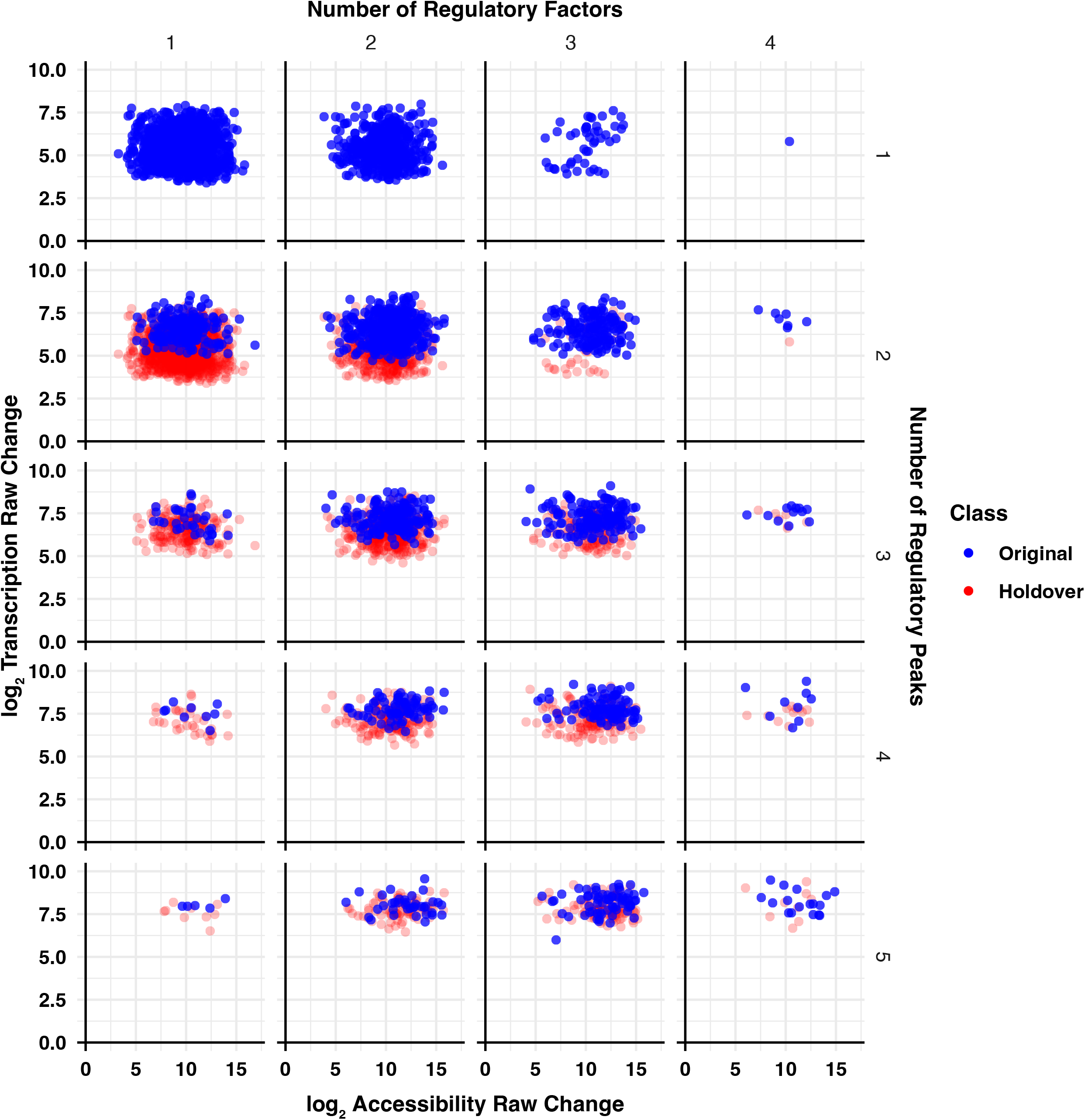
Number of regulatory factors does not influence magnitude of gene transcription changes. All predicted activated target genes are stratified by number of regulatory peaks (rows) and number of regulatory factors (columns). The y-axis indicates the log fold change in transcription for each gene for the time range over which the gene exhibits its greatest activation. Transcription change is plotted against total local accessibility change over the same comparison. Blue points represent the indicated number of regulatory factors and regulatory peaks. Red points represent the genes from the same number of regulatory factors but one fewer regulatory peaks in order to illustrate the effects of an increasing number of peaks. Number of regulatory peaks, and by extension total local accessibility, influences gene transcription. Conversely, the number of regulatory factors does not affect transcription.

**Fig. S8.**
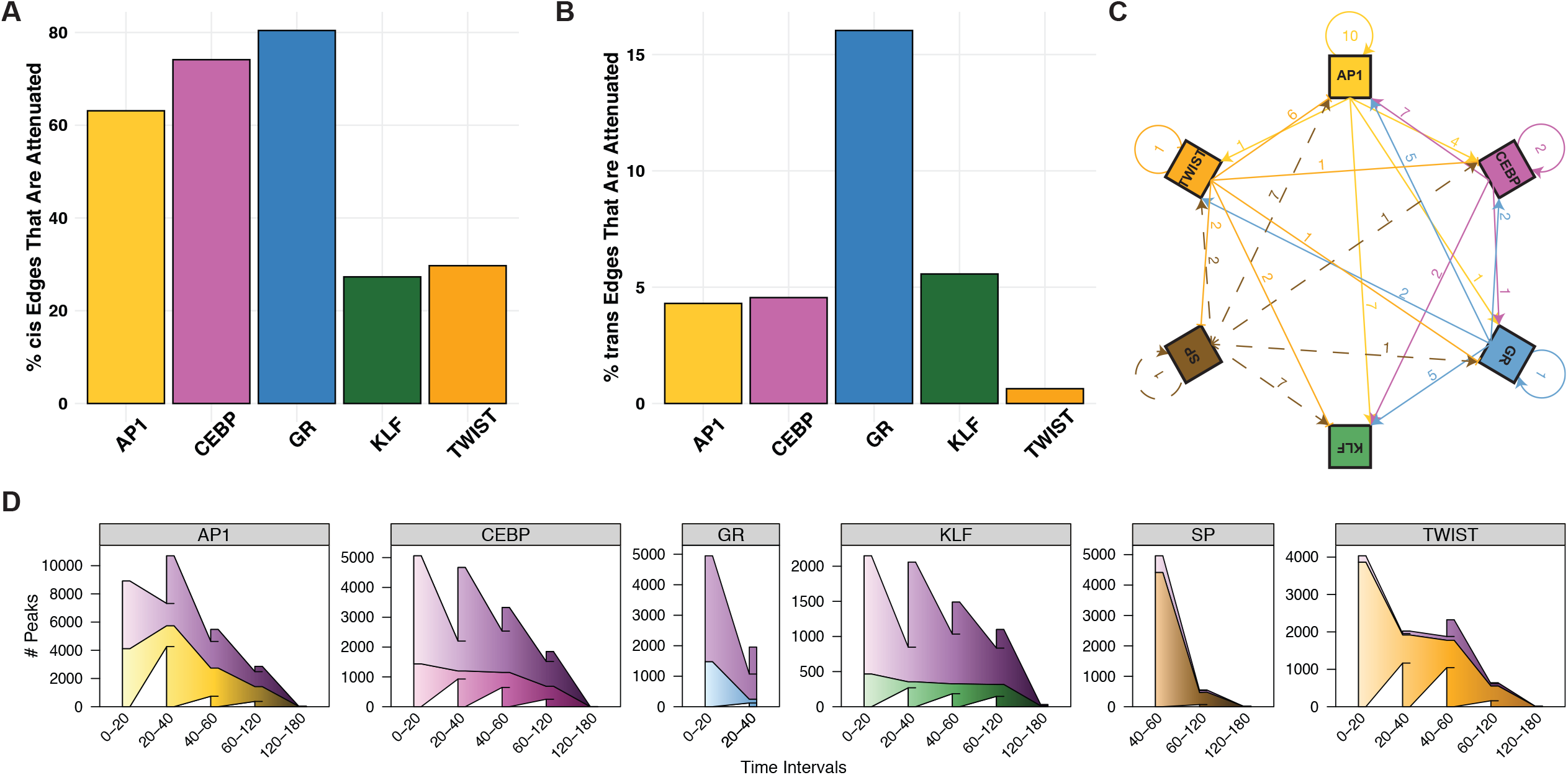
TF families are transiently regulated by other factors, interconnected with one another, and exhibit distinct binding and dissociation kinetics. A) The percentage of each factor’s attenuated *cis*-edges highlights the transient nature of gene expression changes in adipogenesis. Early activators show the highest proportion of attenuated edges, meaning early activation is followed by a return to baseline expression. There are no attenuated SP *cis*-edges, meaning genes with decreased expression downstream of SP dissociation do not return to baseline within the time course. B) The percentage of each factor’s attenuated *trans*-edges indicates that binding is more stable than gene expression changes. GR *trans*-edges are most likely to be attenuated. There are no attenuated SP *trans*-edges. C) TF genes are highly interconnected; arrows represent direct regulatory relationships between factors. Arrows emanate from regulatory factor family and point to target factor family. Solid arrows represent gene activation, blunt-ended arrows represent repression, and dashed arrows represent attenuation. The numbers indicate how many TF family members exhibit the indicated regulatory relationship. For example, the arrow from AP1 to KLF represents the 7 KLF family genes activated by AP1 factors. D) Wedged bar plots quantify the regulatory kinetics across the time course for indicated factors. The x-axis intervals represent the time range in which the specified factor regulates changes in accessibility of the indicated number of peaks (y-axis). Wedges between bars indicate carryover peaks from previous time interval and the outer “wings” represent peaks that are not included in the previous time interval. The top shaded purple wedges represent peaks regulated by multiple factors; bottom wedges represent peaks that are solely regulated by the indicated factor.

**Fig. S9.**
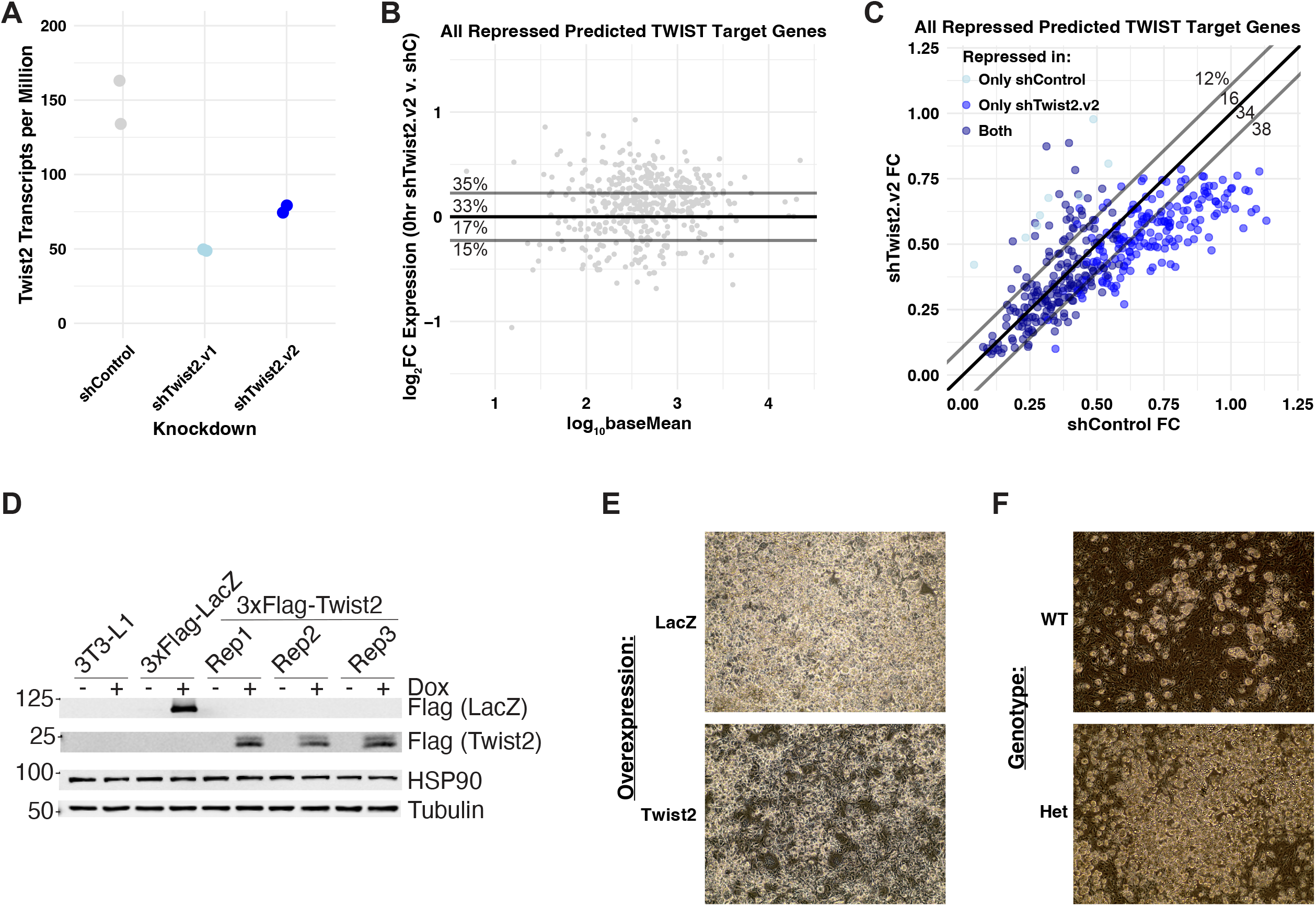
*Twist2* depletion derepresses target genes. A) Baseline expression of predicted TWIST2 target genes repressed in the RNA-seq time course plotted as in Figure 8A). B) Fold change of predicted TWIST2 target genes repressed in the RNA-seq time course plotted as in Figure 8A. C). Similar results with a moderate *Twist2* knockdown further support our hypothesis that TWIST2 acts as a repressor of gene expression and negative regulator preadipocyte differentiation. D) Immunoblotting indicating our tetracycline-inducible 3xFlag-tagged TWIST2 / LacZ constructs work appropriately. HSP90 and Tubulin are used as loading controls. E & F) Images taken at 10x magnification of either (E) 3T3-L1 overexpressing the indicated protein or (F) primary preadipocytes harvested from mice with the indicated genotype.

## Notes

### Competing Interest Statement

The authors have declared no competing interest.

### Summary of Updates

In the revision we compare inferred transcription factor (TF) binding from our network to published ChIP-seq data to molecularly validate the trans-edges in our network. We performed knocked down Twist2 with two separate siRNAs and performed RNA-seq at several time points after induced adipogenesis in the 3T3-L1 system. These results confirmed the inferred cis edges in our network and validated the conclusion that Twist2 is a transcriptional repressor in this context. We also validated the approach to identify genes regulated predominantly by a single TF and the use of compartment modeling to determine which step the TF regulates. We treated a leukemia cell line with dexamethasone to activate the glucocorticoid receptor and confirmed that GR regulates RNA polymerase II pause release. We also performed validation that Twist2 regulates the adipogenic process by depleting and over-expressing Twist2 in the 3T3-L1 system and measuring lipid deposition, which is a measure of adipogenesis. We also performed experiments in Twist2 knockout mice. First, we cultured preadipocytes from Twist2-/+ KO and induced adipogenesis. We found that lipid deposition in the cells was modulated in the heterozygotes. Lastly, we phenotypically characterized Twist2-/+ and Twist2-/- mice. Twist2-/+ have an absence of subcutaneous white adipose tissue and Twist2-/- mice have a near absence of lipid droplets in their brown adipose tissue.

